# Investigating Polyreactivity of CD4+ T Cells to the Intestinal Microbiota

**DOI:** 10.1101/2024.08.15.607895

**Authors:** Ahmed Saadawi, Florian Mair, Esther Rosenwald, Daniel Hoces, Emma Slack, Manfred Kopf

**Author notes:** Current affiliation: Department of Bioengineering, Stanford University, CA, USA.

## Abstract

The symbiotic relationship between host and microbiota plays a pivotal role in training and development of the host’s innate and adaptive immune systems. Antigen-specific recognition of microbiota by T cells enforces tolerance at homeostasis. Conversely, dysbiosis—characterized by alterations in microbiota diversity and abundance—leads to imbalanced T cell responses and triggering of inflammatory and autoimmune diseases. Despite their significance, the identities of immunogenic microbial antigens are still largely enigmatic. Here, we leveraged an in-house developed antigen screening platform, the MCR system ^1^, to delineate CD4+ T cell reactivity against *Akkermansia muciniphila* (AKK) and *Bacteroides thetaiotaomicron* (BT), —two prominent members of the gut microbiota. T-cell hybridomas reactive to AKK and BT bacteria showed polyreactivity to select microbiota-derived peptides in MCR co-cultures. We discovered 13 novel antigenic epitopes from AKK and 14 from BT. Steady-state T cells recognized these epitopes in an MHC-restricted fashion. Ex vivo stimulation of peptide-specific T cells revealed induction of type 1 and type 17 immune responses, albeit with non-overlapping specificities, contrary to MCR system results. Our findings further demonstrated that most identified epitopes are broadly conserved within the given phylum and originate from both membrane and intracellular proteins. Our work showcases the potential of the MCR system for identifying immunogenic microbial epitopes, providing a valuable resource. Additionally, it indicates the existence of mucosal T cells with a tropism toward broadly conserved bacterial epitopes. Overall, our study forms the basis for decoding antigen specificity in immune system-bacterial interactions, with applications in understanding both microbiome and pathogenic bacterial immunity.

## Introduction

The coevolutionary entanglement between microbiota and host has profound implications for host physiology, tissue development, and immune responses against infections ^2–6^. The microbiota exerts pivotal functions in the priming and maturation of the host’s innate and adaptive immune arms. Concurrently, the immune system orchestrates the preservation of essential nutritional attributes within the host-microbiota symbiotic relationship ^7,8^. Nevertheless, the microbiome is linked to the development of inflammatory bowel diseases (IBD), diabetes, obesity, cancer, central nervous system and immune system disorders both in mice and humans ^9–14^. The immune system employs two perception modes of microbiota, comprising coarse-grained and fine-grained recognition. The former encompasses recognition of metabolites and microbe- or pathogen-associated molecular patterns (MAMPs or PAMPs) recognized by the immune system via pattern recognition receptors (PRRs) ^15^. The latter discerns specific antigens derived from microbes at the species or strain level ^16^. Interestingly, even though the gut microbiota is the largest source of intrinsic non-self-antigens present within the intestinal lumen, immune cell subsets are physiologically skewed towards a tolerogenic state under homeostatic conditions ^17–19^. Despite notable efforts in deciphering T cell specificities to microbial antigens, only a few antigens have been identified and validated. This intricate balance underscores the complexity of mucosal immune responses.

It is well established that various gut microbial species can drive the differentiation of different flavors of CD4+ T helper cell subsets in an antigen-dependent manner. Reportedly, colonization of germ-free (GF) mice with the commensal segmented filamentous bacteria (SFB) potently induced the accumulation of T helper 17 (T_H_17) cells ^20^. Subsequent analysis showed that most T cell receptors (TCRs) of the intestinal T_H_17 cells induced by SFB recognized two nine–amino acid peptides, suggesting that the T cell response was directed toward select dominant peptides^21^. Members of the genus Clostridium, particularly clusters IV and XIV, have been demonstrated to promote peripheral regulatory T cell (pT_regs_) accumulation ^22,23^. Additionally, the pathobiont *Helicobacter hepaticus* has been shown to induce the differentiation of pT_regs_ and CD4^+^ T follicular helper cells (T_FH_) ^24^. Moreover, *Bacteroides thetaiotaomicron* (BT) and *Akkermansia muciniphila* (AKK) have been described to induce a distinct population of Rorγ^+^ T_regs_ in the mouse colon, which controls inflammation and maintains mucosal homeostasis ^25–29^.

Importantly, T cells specific for commensals can adopt different fates depending on the context and the antigen recognized. Evidently, T cells specific for AKK-derived epitopes predominantly adopted a T_FH_ phenotype in GF mono-colonization settings ^30^. However, in conventional mice, the T cell response to AKK was a mixture of multiple subsets (T_H_1, T_H_17, T_regs_, and T_FH_) ^30^. In another study, T cells reactive to BT-derived peptide manifested an interleukin 10 (IL-10) cytokine profile in healthy individuals, which shifted to an IL-17A profile in Crohn’s disease patients, demonstrating the context-dependent modulation of T cell fate resulting from host-microbiome interactions ^31^.

BT and AKK are two of the most abundant and most intensively studied gut bacteria. They are strictly anaerobic, gram-negative bacteria colonizing the human intestinal lumen and mucosal layer and belong to the Bacteroidetes and Verrucomicrobia phyla of bacteria, respectively. BT degrades dietary fiber polysaccharides and host glycans, whereas AKK functions as a mucin-degrader, thereby influencing mucus layer thickness ^32–34^. Combined, they often constitute >10% of total intestinal bacteria population density in humans ^35,36^. BT and AKK have been associated with a plethora of biological functions. On the one hand, BT facilitates nutrient absorption and exerts protective functions by producing outer membrane vesicles. These vesicles can cross the intestinal epithelial barrier to help maintain intestinal homeostasis through the reciprocal crosstalk with both the host’s epithelial and immune cells ^37,38^. In contrast, BT has been shown to aggravate metabolic disorders by regulating lipid metabolism ^39,40^, and to trigger colitis in genetically susceptible mice ^41^. On the other hand, AKK is densely populated in the intestinal epithelium and its strategic position at the mucosal interface between the lumen and host cells renders it a sentinel for gut permeability, thus providing increased protection against pathogen infections ^42^. Several lines of evidence have pleaded in favor of the beneficial roles of *AKK*. It was shown to reduce obesity ^43,44^; to improve glucose homeostasis and ameliorate metabolic disease ^45–47^; and to enhance the efficacy of anti PD-1–based immunotherapy against tumors ^48–50^.

Previous research has identified very few BT- and AKK-derived epitopes in the context of major histocompatibility complex (MHC) class II. However, technical limitations have hampered efforts to identify bacteria-specific T cells systematically ^26,30,51–53^. Here, we utilized the MCR system, — a sensitive, unbiased and genome-wide platform for the systematic identification of BT- and AKK-derived epitopes recognized by CD4+ T cells. The MCR platform allows for high-sensitivity discovery of MHC class II restricted epitopes. It is based on screening of peptide libraries using an NFAT-fluorescence timer reporter cell line (i.e., thymoma) that is transduced with a retrovirus encoding an MHC-TCR (MCR) hybrid molecule. The MCR consists of the transmembrane and intracellular part of the TCR (α) and (β) chains fused to the extracellular portion of the of MHCII (α) and (β) chain that is genetically tethered to a unique peptide **(Fig. 1a**). MCR-positive cells could be isolated based on surface expression of CD3e and/or MHC II. Co-culture of specific T cells with the library allows the monitoring of specific TCR-peptide-MCR interaction by switching on the expression of the fluorescent reporter in the cell presenting the cognate target peptide ^1^. The expression of the fluorescence timer, with blue indicating recent NFAT activation (blue-FT) and red indicating past activation (red-FT), enhances signal resolution.

**Fig. 1:**
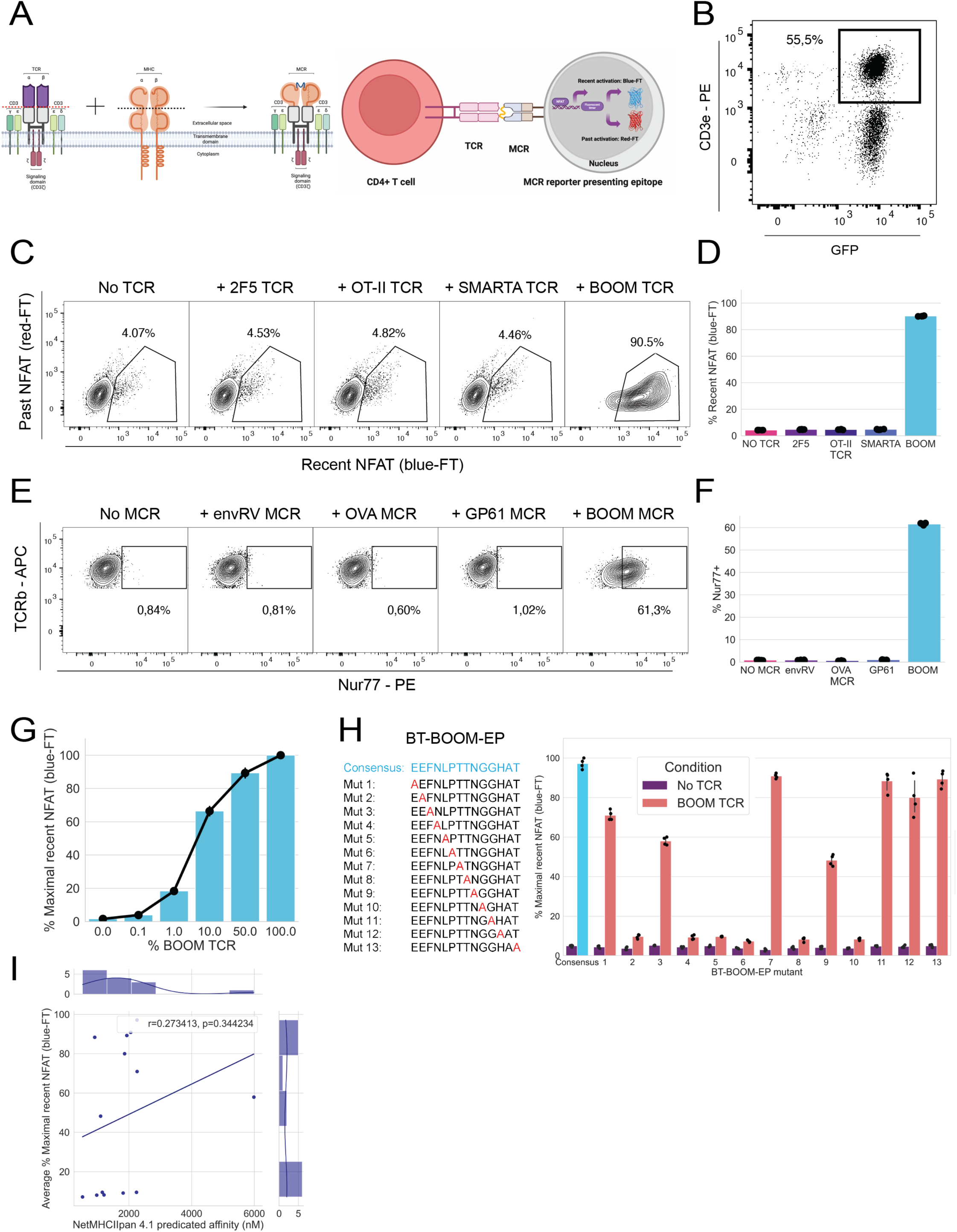
MCR-based model system for identification of microbiota-derived CD4+T cell specific epitopes. a, Cartoon representation of the MCR and its interaction with peptide-specific TCR. Figure was created with BioRender.com **b**, Surface expression of the MCR carrying Bacteroides thetaiotaomicron BθOM epitope (BT- BθOM-EP^+^). 16.2X reporter cell line is transduced with BT-BθOM-EP^+^ MCR plasmid and stained for CD3e. 16.2X reporter cells constitutively express GFP. **c**, Specificity of detection of peptide-specific T cells. BT-BθOM- EP^+^ MCR+ reporter cells are left untreated or co-cultured with different TCRs in a 1:5 ratio and NFAT activation is measured after 16 h. 2F5 TCR, OT-II TCR, SMARTA TCR and BΘOM TCR are specific for envRV, OVA 323-339, GP61 and BT-BθOM-EP^+^ peptides, respectively. **d,** Quantification of **c**, (*n* = 25 co-cultures). **e,** MCR- based induction of Nur77 in T cell hybridoma. BθOM TCR is left untreated or co-cultured with different MCRs in a 5:1 ratio and Nur77 upregulation is measured after 4 h. **f,** Quantification of **e**, (*n* = 25 co-cultures). **g,** Minimal frequency of peptide-specific T cells able to trigger robust NFAT activation in MCR+ reporter cells. The graph depicts percentage of maximal NFAT activation of BT-BθOM-EP^+^ MCR+ reporter cells as a function of the percentage of BΘOM TCR (determined by FACS), after 12 h of co-culture (*n* = 24 co-cultures). **h,** Scanning alanine mutagenesis of BT-BθOM-EP. Systematic alanine substitution of every non-alanine amino acid in BT- BθOM-EP (n= 13 mutants). Consensus BT-BθOM-EP is highlighted in cyan (left panel). MCR+ reporter cells expressing BT-BθOM-EP mutants or consensus are left untreated or co-cultured with BΘOM TCR in a 1:5 ratio (*n* = 112 co-cultures). Percentage of maximal NFAT activation is measured after 16 h of co-culture (right panel). **i,** Correlation between NetMHCIIpan 4.1 predictions and experimental NFAT activation measurements. Pearson’s correlation between NetMHCIIpan 4.1 predicted affinity and experimentally measured NFAT activation of BT- BθOM-EP mutants (averaged) after co-culture with BθOM TCR. Data are representative of two (**c,e,g,h**), independent experiments.

By generating T cell hybridomas reactive to BT and AKK bacteria, conducting iterative enrichment of target epitopes, and coupling next-generation sequencing (NGS) with *in vivo* mouse immunizations, we identified several BT- and AKK-derived immunogenic epitopes that induced IL-17A and IFN-γ responses and are conserved in several members of the Bacteroidetes and Verrucomicrobia phyla.

## Results

### Setting up the MCR system for identification of microbial antigens

As a prelude to our experiments, we set out to assess the specificity and sensitivity of the MCR system for bacteria-derived epitopes by employing a previously identified BT-derived epitope (i.e., BθOM peptide) and its cognate TCR expressed by a hybridoma (i.e., BθOM-TCR_H_) ^26^. To that end, we expressed the BθOM peptide in MCR reporter cells (BT-BθOM-EP^+^ MCR) and co-cultured them with CD4^+^ T cell hybridomas carrying different TCRs including BθOM-TCR_H_ **(Fig. 1b)**. As expected, the NFAT signal was robustly and specifically upregulated in BT-BθOM-EP^+^ MCR co-cultured with BθOM- BθOM-TCR_H_ but not with hybridomas carrying other TCRs **(Fig. 1c,d)**. To further examine whether BT-BθOM-EP^+^ MCR would induce signaling in BθOM-TCR_H_, we co-cultured BθOM-TCR_H_ with different MCRs. As expected, Nur77, a downstream signaling molecule whose expression is rapidly activated upon TCR signaling, was upregulated in the co-culture of BθOM-TCR with BT-BθOM-EP^+^ MCR, but not with other MCRs, indicating specific reciprocal recognition **(Fig. 1e,f)**. Next, we sought to determine the sensitivity of detection. To tackle this, we initially probed the minimal frequency of BθOM-TCR_H_ required for induction of robust NFAT signal. Accordingly, we mixed escalating percentages of cognate BθOM- TCR_H_ with another non-cognate T cell hybridoma (i.e., BT-H1) to BT-BθOM-EP^+^ MCR. Our results showed that the presence of only 1% BθOM-TCR_H_ in the co-culture mixture stimulated 18% of maximal NFAT activation in BT-BθOM-EP^+^ MCR cells **(Fig. 1g)**. Furthermore, we conducted systematic alanine mutagenesis of the BθOM peptide, generating 13 mutants. Each mutant was expressed in MCR reporter cells and individually co-cultured with the BθOM-TCR_H_. Interestingly, certain mutations (numbers: 2, 4, 5, 6, 8, 10) completely abrogated the binding of BθOM TCR with mutant BT-BθOM-EP^+^ MCR, evidenced by lack of NFAT signal, likely due to disruption of peptide-MHC II anchor sites and/or TCR-peptide MHC II specific contact points. Additionally, other mutations (numbers: 1, 3, 7, 9, 11, 12, 13) reduced the NFAT signal by 20 – 50 %, compared to consensus BT- BθOM-EP^+^ MCR **(Fig. 1h)**. Finally, we utilized netMHCIIpan ^54,55^ to predict binding affinity of the BθOM peptide mutations to mouse MHC II (I-Ab) molecule and correlated these predictions with experimental NFAT signal measurements derived from MCR co-cultures. Notably, Pearson’s correlation coefficient was low (r = 0.27), indicating that netMHCIIpan’s predictions did not fully capture the experimental data **(Fig. 1i)**. Taken together, these data show that the MCR system can produce specific and sensitive signals for a bacterially-derived peptide, indicating promise for investigating more complex peptide mixtures.

### Generation of BT and AKK reactive T cell hybridomas and MCR libraries

We aimed to identify novel peptide epitopes from BT and AKK bacteria. Our experimental design **(outlined in Fig. 2)** consisted of two primary compartments that converged at the screening phase. First, we constructed genome-wide MCR libraries of BT and AKK bacteria **(Supplementary** Fig. 1a). These libraries covered the entire bacterial genome around 600 times, sufficient for comprehensive representation of DNA-encoded peptides and enabling deep exploration of TCR specificity **(Supplementary** Fig. 1d). In parallel, we immunized GF and specific pathogen-free (SPF) mice intratracheally (i.t.) with heat-killed (HK) bacteria to generate a high number of bacteria-reactive CD4^+^ T cells. We opted for harvesting CD4^+^ T cells from the lungs of i.t. immunized mice over isolating CD4^+^ T cells from the gut due to the lower percentage of viable BT and AKK-reactive T cells that can typically be isolated from the gut, linked to high concentrations of toxic metabolites in digesta and long digestion steps required for the lamina propria. After sorting activated T cells from the immunized mice, we fused them with an immortal thymoma line, generating 8 BT-reactive and 9 AKK-reactive T cell hybridomas derived from a mix of GF and SPF mouse immunizations **(Supplementary** Fig. 1b,c**,e)**. Of note, activated T cells showed reduced capacity to generate T cell hybridomas compared to naïve non-activated T cells, yielding three times fewer hybridomas in our experiments, and explaining the low number of microbiota-reactive T cell hybridomas (data not shown). Importantly, the generated T-cell hybridomas upregulated CD69 after co-culture with bone marrow derived dendritic cells (BMDCs) pulsed with bacteria, confirming their cognate reactivity toward the immunizing bacteria (Supplementary Fig. 2). However, we observed cross-reactivity of the T cell hybridomas to non-immunizing bacteria— BT-reactive hybridomas respond to AKK bacteria, and similarly, AKK-reactive hybridomas respond to BT bacteria (Supplementary Fig. 2). During the screening phase, one or more T cell hybridomas were co-cultured with the cognate MCR library, and successive enrichment of target peptides was performed **(Supplementary** Fig. 3a). Single MCR+ reporter cells expressing the blue-reporter from the final round of enrichment were sorted, expanded, and recalled with the same T cell hybridoma(s) used initially for screening **(Supplementary** Fig. 3b). Sequencing of reactive reporter cells unveiled the identity of the recognized peptides. To systematically identify peptides recognized in the final round of enrichment, we additionally conducted amplicon NGS on the peptide-encoding fragment of the MCR plasmid **(Supplementary** Fig. 3c,d). NGS data demonstrated that the most abundant peptide constituted ≤ 50% of the total sequencing reads and was incorporated in overlapping amino acid (AA) configurations **(Supplementary** Fig. 3e,g**,h)**. Moreover, the length of DNA-encoded peptides remained consistent across all rounds of enrichment **(Supplementary** Fig. 3f). Collectively, these data indicate the successful enrichment of target peptides recognized by microbiota-reactive T cells using the MCR system.

**Fig. 2:**
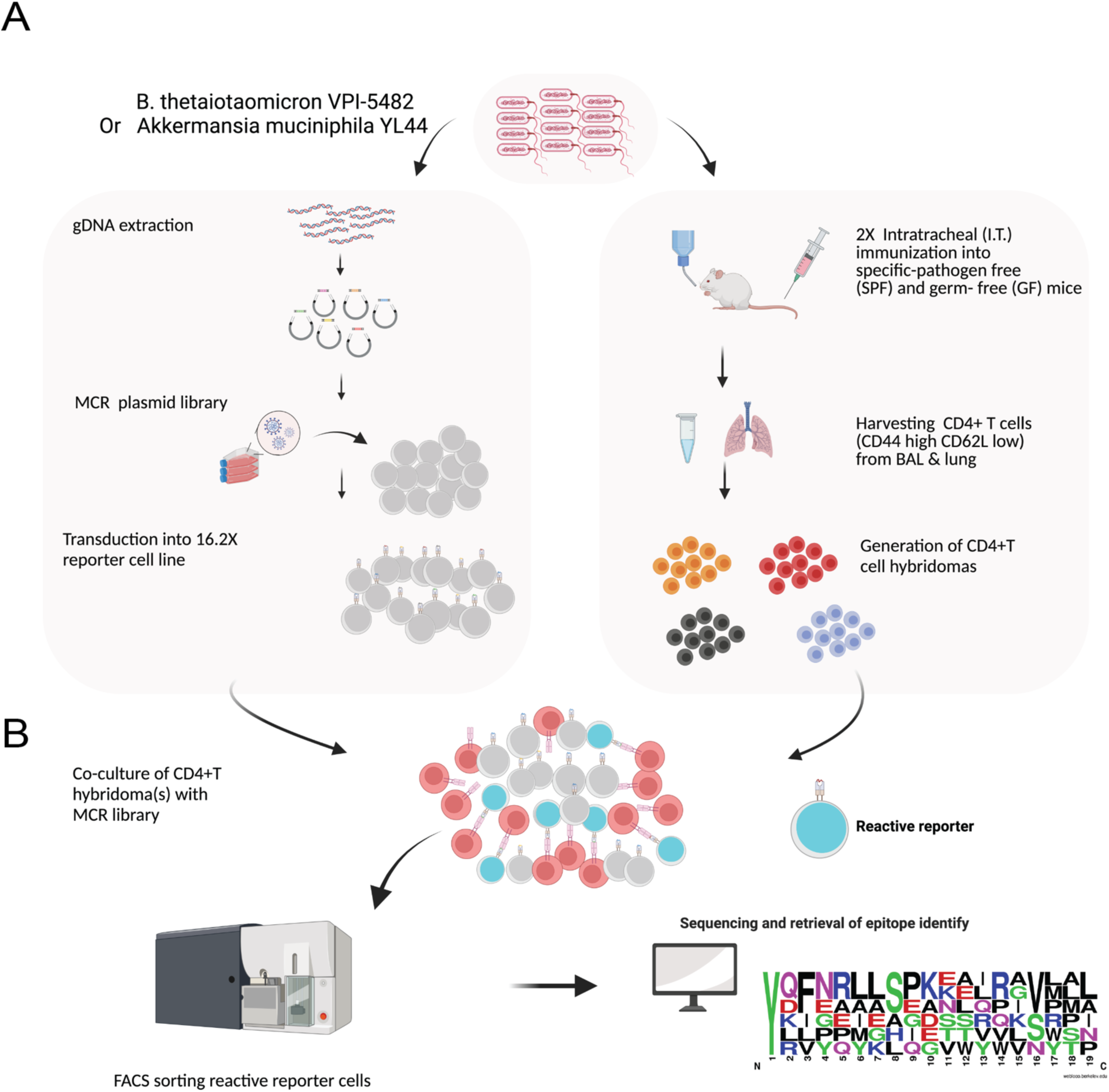
Methodology for generation of MCR libraries and CD4+T cell hybridomas. a, Bacterial genomic DNA is sheared into 150 base pairs (bp), on average, ligated to 100 bp linkers, and cloned into the MCR plasmid. MCR negative 16.2X reporter cells are transduced with MCR plasmids and MCR+ cells are FACs sorted, thereby constituting an MCR library. In parallel, specific-pathogen-free (SPF) and germ-free (GF) mice are immunized twice intratracheally (i.t.) over the period of two weeks. Bronchoalveolar lavage (BAL) and lung are harvested and activated CD4+T cells are FACs sorted and fused to α^−^β^−^ (TCR negative) BW5147 thymoma to generate T cell hybridomas. CD4+T cells hybridomas are derived from both SPF and GF mice. **b,** Bacteria-specific MCR library reporter cells are co-cultured with cognate T cell hybridoma (s) of interest and activated MCR+ reporter cells are FACs sorted, expanded for one week and co-cultured again with the same T cell hybridoma(s). This step is repeated for 4-6 rounds to enrich for target peptides in the library. Finally, at the last round of iterative enrichment, single activated or population of reactive MCR+ reporter cells are sorted and expanded, where the peptide-encoding fragment of the MCR plasmid is PCR amplified and subjected to Sanger sequencing or amplicon next-generation sequencing (NGS), respectively, to reveal encoded peptide sequence. Figure was created with BioRender.com

### Profiling T cell reactivity to microbiota-derived peptides

Having successfully completed screenings for all BT- and AKK-reactive T-cell hybridomas, we selected the most enriched candidate peptides and compiled them into “mini” libraries. The BT “mini” library encompassed 30 candidate peptides **(Table 1)**, whereas the AKK “mini” library comprised 19 candidate peptides **(Table 2)**. These peptides exhibited significant length variation, ranging from 17 to 48 amino acids (AAs) for BT and 16 to 55 AAs for AKK, with average lengths of 29 AAs and 33 AAs, respectively. Notably, peptides binding to MHC class II molecules typically range from 11 to 30 residues ^56^. Derived from both membrane and intracellular proteins, these peptides reflected a diverse range of cellular localizations.

**Table 1:**
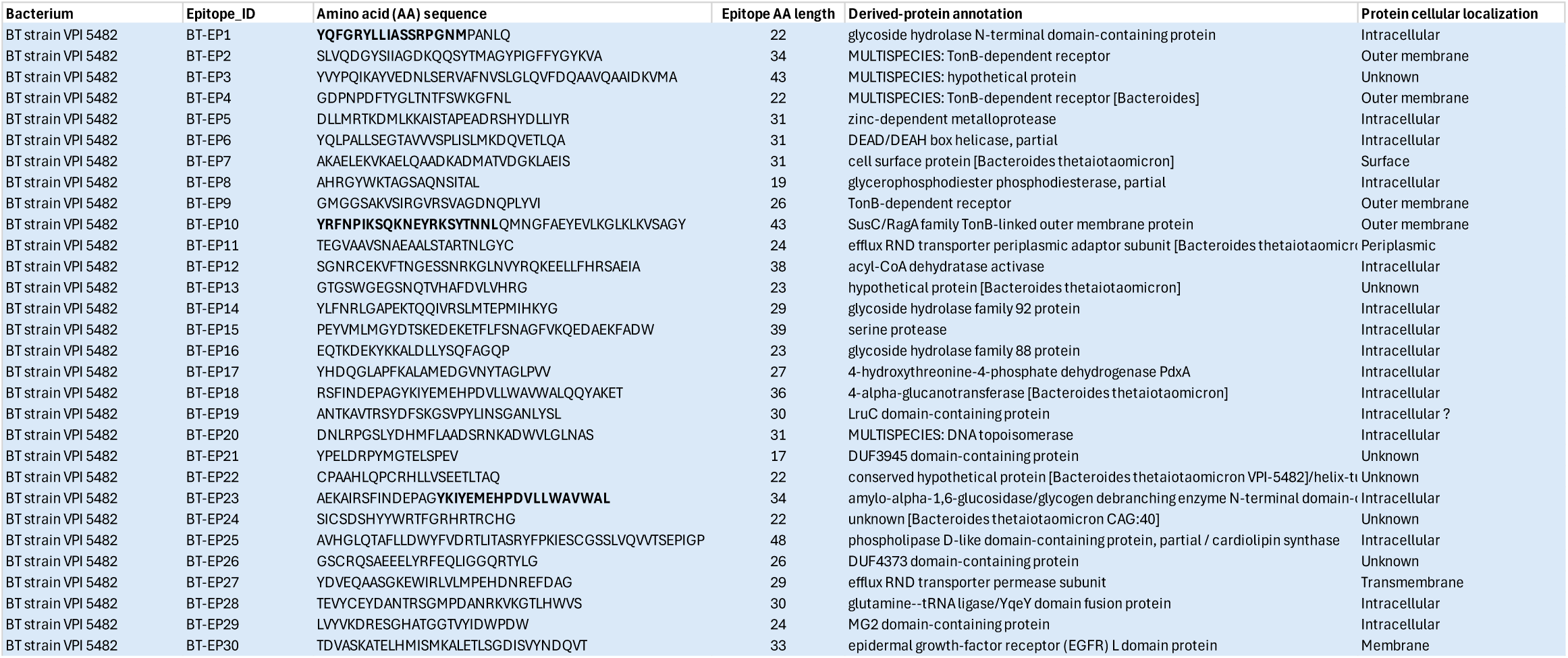
Bacteroides thetaiotaomicron (BT) peptides’ information.

**Table 2:**
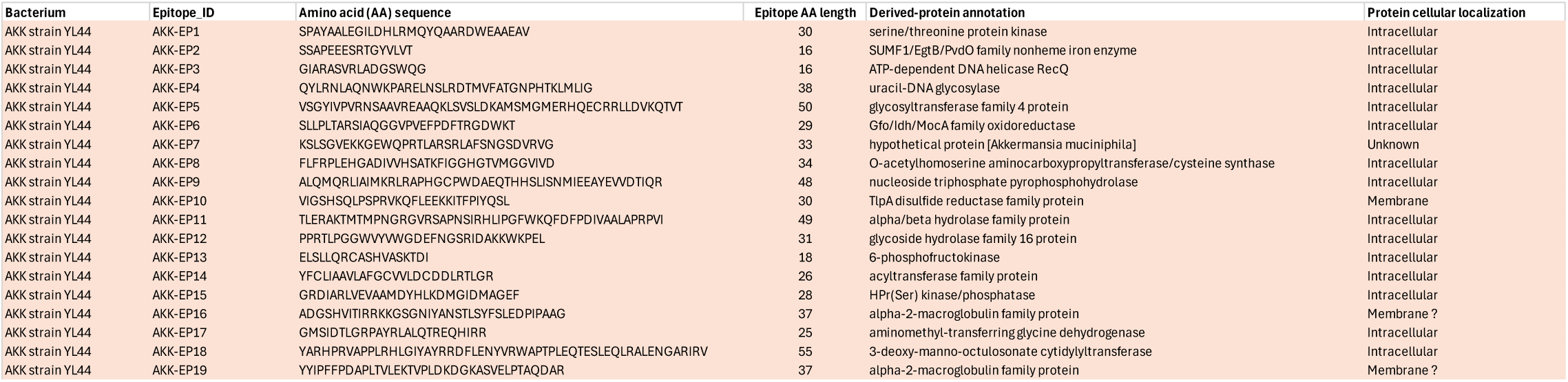
Akkermansia muciniphila (AKK) peptides’ information.

To discern patterns of reactivity, we freshly transduced these peptides individually into MCR-negative reporter cells and co-cultured them with BT- and AKK-reactive T-cell hybridomas. Intriguingly, BT-reactive T-cell hybridomas reproducibly recognized select peptides from both BT and AKK bacteria, eliciting differential NFAT signal intensities suggestive of varied TCR binding strengths (**Fig. 3)**. Specifically, AKK MCRs displayed quantitatively higher signal intensities. Overall, BT-reactive T-cell hybridomas reacted with 14 out of 30 BT-derived peptides and 13 out of 19 AKK-derived peptides. Similarly, AKK-reactive T cell hybridomas induced NFAT signaling in both AKK- and BT- derived peptides, the same MCR^+^ reporter cells that had shown responses with BT-reactive T cell hybridomas (**Fig. 3)**. Crucially, neither BT- nor AKK-reactive T cell hybridomas exhibited reactivity towards peptides from mammalian, viral, or other bacterial origins, underscoring selective recognition of microbiota-derived peptides (**Fig. 3)**. Comparatively, control T cell hybridomas from naïve, non-immunized mice demonstrated no reactivity to the microbiota-derived peptides (**Fig. 3)**. It is noteworthy that spontaneous loss of TCR expression on BT- or AKK-reactive T cell hybridomas completely abolished NFAT signaling in responsive MCRs, confirming the exclusive role of TCR-MCR binding and excluding the contribution of other accessory molecules on activated T cells in the observed responses (data not shown). In summary, our findings suggest a bias towards recognition of broadly conserved peptides by bacteria-reactive CD4^+^ T cells *in vivo,* as depicted in MCR co-cultures.

**Fig. 3:**
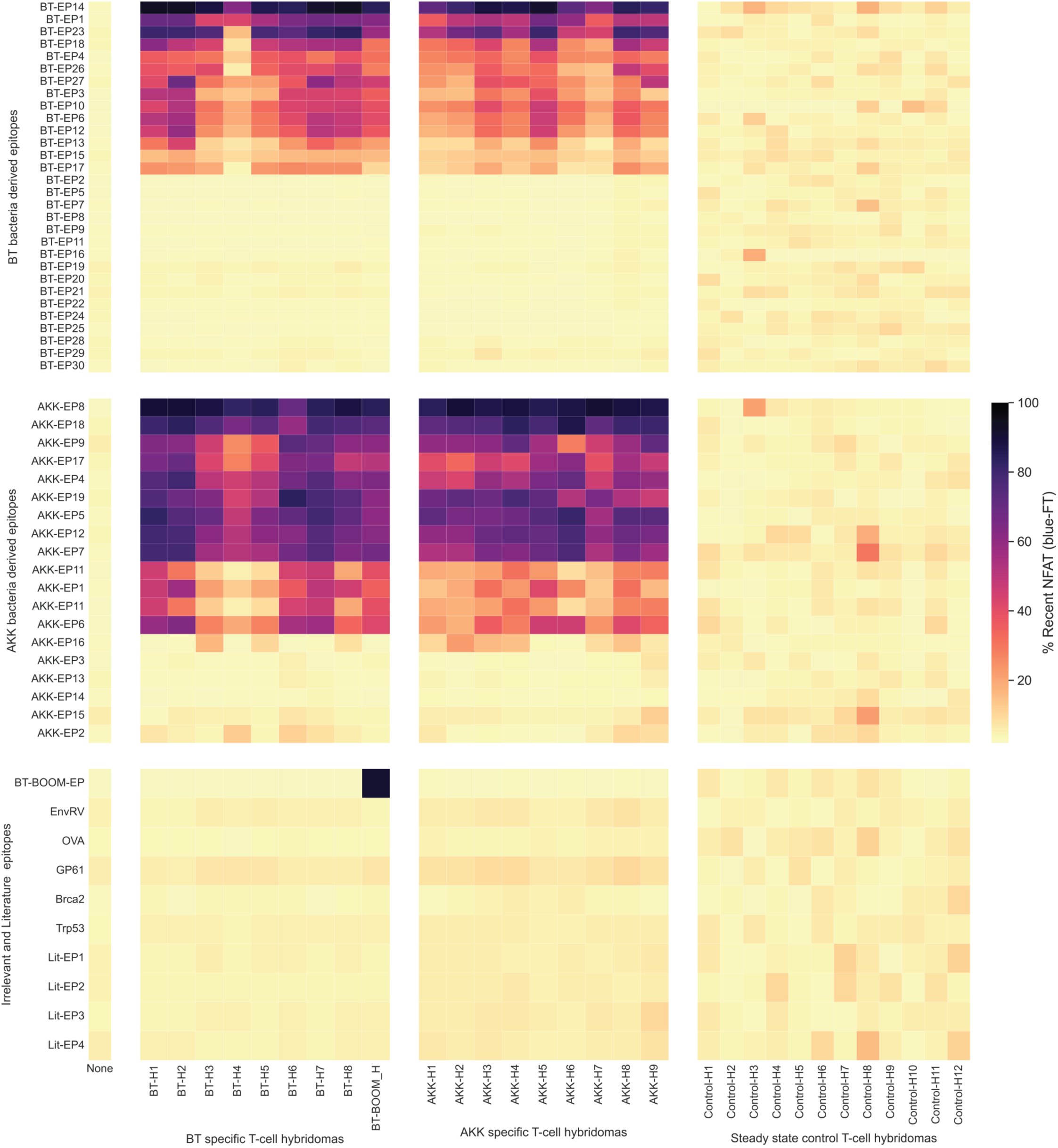
Widespread poly-reactivity of microbiota specific CD4+T cell hybridomas in MCR co-cultures. Mapping of T cell epitope specificity. Shown is a 3x3 matrix of pools of T cell hybridomas and epitopes. The first two pools of T cell hybridomas are derived from “activated” T cells mounted against Bacteroides thetaiotaomicron (BT) and Akkermansia muciniphila (AKK), respectively. The third pool of T cell hybridomas is derived from “steady state” T cells harvested from naïve nonimmunized SPF mice, — i.e., CD62L high and CD44 medium/low phenotype (right column panel). BT-H-# indicates hybridomas derived from mice immunized with BT (left column panel), and AKK-H-# indicates hybridomas derived from mice immunized with AKK (middle column panel). BT-H1 and BT-H2 are derived from GF immunized mice, while all other BT-H# hybridomas are derived from SPF immunized mice. Likewise, AKK-H1, AKK-H2 and AKK-H3 are derived from GF immunized mice, while all other AKK-H# hybridomas are derived from SPF immunized mice. Similarly, the first two pools of epitopes comprise BT (upper row panel) and AKK (middle row panel) candidate epitopes, respectively. BT and AKK epitope pools are composed of highly abundant inframe epitope-encoding DNA sequences from BT-reactive and AKK-reactive T cell hybridomas’ screenings, respectively. BT-EP# indicates epitopes derived from screenings of BT-reactive hybridomas, and AKK-EP-# indicates epitopes derived from screenings of AKK- reactive hybridomas. Third pool designated as “Irrelevant and Literature epitopes,” includes epitopes derived from viral, mammalian, and other bacterial sources (lower row panel). Lit-EP-# indicates epitopes derived from previously reported microbial epitopes. EnvRV and GP61 are viral epitopes. Brca2 and Trp53 are mammalian epitopes. Each T cell hybridoma is co-cultured with every epitope individually and NFAT activation was measured after 16 hours. Colorbar represents percentage of recent NFAT activation. Two independent efforts of cloning, transduction and co-culture are conducted from scratch. The data are an average of two independent experiments per transduction and co-culture effort.

We then investigated whether T cells of irrelevant specificities would recognize our identified microbiota-derived epitopes. To achieve this, we co-cultured T-cell hybridomas specific for influenza virus and the mammalian murine mammary cancer cell line EO771 with BT, AKK, and irrelevant epitopes expressed in the MCR system **(Supplementary** Fig. 4). Surprisingly, some of these viral and mammalian-specific T cell hybridomas (VMTs) recognized select peptides from both BT and AKK bacteria—the same epitopes that had elicited responses with BT- and AKK-reactive T cell hybridomas. However, unlike the BT- and AKK-reactive T cell hybridomas, where one hybridoma recognized all selected epitopes, some VMTs recognized fewer microbiota-derived epitopes. Additionally, the NFAT signal intensity varied among the different VMTs, suggesting varied TCR binding strengths. Notably, the VMTs exhibited no reactivity towards peptides from other microbial sources. Furthermore, we observed that OT-II and 2F5 T cell hybridomas, specific for OVA and envRV peptides respectively, recognized these select MCR epitopes in addition to their cognate peptides. Collectively, our findings indicate that the MCR-fused epitopes we identified can recognize a diverse array of T-cell receptors (TCRs) with varying specificities. This broad recognition pattern is reminiscent of the behavior observed in superantigens, suggesting a potential for these epitopes to engage in widespread immune activation.

Microbiota-specific T cells were shown to exist in the periphery at steady state ^57^. In addition, the trafficking of microbial antigens to the thymus by intestinal CX3CR1^+^ DCs drives expansion of microbiota-specific T cells ^58^. Accordingly, we aimed to examine whether primary T cells from the spleen and thymus of naive mice would recognize any of our microbiota-derived peptides. We co-cultured CD4^+^ T cells obtained from spleen and thymus of C57BL/6 mice maintained either in a germ-free or (GF), or specific pathogen-free (SPF) environment, and the spleen of SPF BALB/c mice with BT- and AKK-derived MCR reporter cells **(Supplementary** Fig. 5a). We observed that both splenic and thymic CD4^+^ T cells from C57BL/6 mice recognized select BT- and AKK bacteria derived peptides, — the same reactive peptides with BT- and AKK- specific T cell hybridomas. To delineate which CD4^+^ T cell subset is driving this response, we co-cultured T_reg_ or conventional T cells (T_con_), sorted from SPF DEREG mice, with BT- and AKK- MCRs. Both T_reg_ and T_con_ recognized the select peptides, suggesting a balanced T_reg_ / T_con_ immune response. Interestingly, GF splenic CD4^+^ T cells induced NFAT signaling in some MCRs. However, the NFAT signal was 50% reduced compared to coculture with SPF splenic cells, indicating enrichment of microbiota-reactive T cells in SPF vs GF mice. Reportedly, some bacteria restimulate intestinal CD4^+^ T cells from GF mice ^29^. Expectedly, CD4^+^T cells from BALB/c mice elicited no reactivity to any of the MCRs, confirming MHC restriction basis of T cell recognition. Collectively, our data indicate that different subsets of steady-state T cells, possibly representing a spectrum of affinities, recognize our microbiota-derived peptides in an MHC-restricted manner.

To pinpoint the core binding domain of some reactive microbiota-derived peptides, we transduced extending and overlapping fragments of BT-EP1 and BT-EP10, respectively, into MCR negative reporter cells and co-cultured them with BT- and AKK-reactive T cell hybridomas **(Supplementary** Fig. 5b). The 17-amino acid fragment of BT-EP1 displayed similar NFAT response as the full-length BT-EP1 (22 AAs) with all different T cell hybridomas tested, revealing it to be the core binding site. However, shorter fragments of BT-EP1 (14, 15, and 16 AAs) also showed some reactivity with select T cell hybridomas. Similarly, the core binding site of BT-EP10 (43 AAs) was trimmed down to be 21 AAs **(Supplementary** Fig. 5c).

We then wondered whether reactive microbiota-derived peptides possess a conserved binding motif. Using Clustal Omega for multiple sequence alignment (MSA), our analysis unraveled no shared common motifs among the peptides, suggesting their distinct and diverse nature **(Supplementary** Fig. 6a,b). However, we observed a tyrosine (Y) amino acid conservation at position one in most BT peptides and some AKK peptides. Tyrosine residues (often called anchor residues) are known to play a crucial role in the binding of peptides to to MHC class II molecules ^59^.

### BT- and AKK-reactive T cell hybridomas are not activated by identified bacterial synthetic peptides

Next, we aimed to probe whether the bacteria-reactive T-cell hybridomas would recognize the identified epitopes in a system other than the MCR. In a confirmatory experiment, we co-cultured BT-reactive T-cell hybridomas with some epitopes derived from both BT and AKK bacteria expressed in MCR+ cells (**Supplementary** Fig. 7a). As expected, BT-reactive T cell hybridomas exhibited the same pattern of reactivity toward select microbial epitopes as observed previously (refer to Fig. 3). Additionally, we assessed levels of Nur77 in MCR+ cells, revealing a consistent and concomitant pattern of NFAT and Nur77 upregulation (**Supplementary** Fig. 7b). We then examined whether the MCR+ cells would induce Nur77 in the cognate T cell hybridomas. As expected, the BT-BθOM-EP^+^ MCR+ cells induced Nur77 increase in the BθOM T cell hybridoma (**Supplementary** Fig. 7c).

Intriguingly, however, neither BT nor AKK MCR+ cells induced Nur77 expression in BT-reactive T cell hybridomas (**Supplementary** Fig. 7c). Furthermore, fusion of epitopes to MHC II and expression in BMDCs induced no Nur77 upregulation in BT-reactive T cell hybridomas (data not shown). We then co-cultured BT- reactive T-cell hybridomas with BMDCs pulsed with the microbiota-derived peptides (**Supplementary** Fig. 7d). Surprisingly, we found that the BT-reactive T cell hybridomas were not activated by the synthetic peptides, evidenced by a lack of Nur77. In contrast, the BθOM T cell hybridoma recognized their cognate BθOM peptide.

We then questioned whether the bacteria peptides bind to the MHC with low affinity. Therefore, we titrated the concentration of peptides, spanning a 30-fold range from 1 to 300 μg/ml of peptide. However, BT-reactive T cell hybridomas did not recognize the peptides (data not shown). We then wondered whether the TCR signaling machinery was intact in the T cell hybridomas. Accordingly, we stimulated the BT-reactive T cell hybridomas with anti-CD3e antibodies, revealing induction of Nur77 (**Supplementary** Fig. 7c,d).

In summary, our data indicate that the bacteria-reactive T cell hybridomas do not recognize the identified BT and AKK peptides in a system other than the MCR, suggesting a structural mechanism underpinning the contrasting results observed between the MCR-fused and synthetic peptide epitopes.

We then wondered whether the identified bacterial peptides act as antagonists for TCR signaling. To test this hypothesis, we co-cultured the BθOM T cell hybridoma with the BθOM peptide in the presence or absence of other bacteria-derived or irrelevant peptides. As expected, the BθOM peptide, either expressed in the MCR or pulsed in BMDCs, robustly induced Nur77 upregulation in the BθOM T cell hybridoma, while all other peptides had no effect (**Supplementary** Fig. 8a,b). Our results further demonstrated that bacteria-derived peptides, whether expressed in the MCR or pulsed in BMDCs, had no significant effect on Nur77 levels when the BθOM T cell hybridoma was co-cultured with the BθOM peptide. Together, these data indicate that the identified microbiota-derived peptides do not act as antagonists for TCR signaling.

### BT- and AKK- derived peptides elicit a T-cell response *in vivo* after immunization

Microbiota-specific T cells can adopt various effector phenotypes, predominantly T_H_1, T_H_17, and T_regs_, with associated secretion of IFNγ, IL-17A, and IL-10 cytokines, respectively. Prior research has shown that gut T cells are specific to bacterial strains at the species level, can discern between strains of the same species, and secrete IFNγ and IL-17A upon ex vivo antigen encounter ^16^. To investigate the *in vivo* immunogenicity and functional characteristics of T cell responses to identified microbiota-derived peptides, we selected six representative peptides for BT and three for AKK from the reactive MCRs. These peptides manifested similarly high levels of NFAT signal and encompassed both surface and intracellular proteins, with functions in glycoside hydrolysis, binding and transportation of siderophores, and cytochrome biogenesis and maturation, among others. We immunized C57BL/6J SPF mice subcutaneously with each peptide individually and assayed T cell recall responses to an array of peptides from different sources using IL-17A and IFN-γ ELISAs. We found that five out of six BT peptides and two out of three AKK peptides exhibited immunogenicity, as indicated by IL-17A secretion **(Fig. 4a,b)**. However, the recall response was heterogeneous and varied in magnitude. Specifically, the immunogenic peptides manifested a diverse spectrum of IL-17A secretion, spanning a range from 2 to 30 ng/ml. Strikingly, the T-cell response was exclusively directed to the cognate immunizing peptide and did not display the polyreactivity observed in MCR co-cultures. IFN-γ cytokine secretion showed similar patterns of immunogenicity and lack of polyreactivity (data now shown). This is likely explained by the fact that the MCR system captures MHC-TCR interaction only, while induction of an i*n vivo* response relies on additional signals (specifically co-stimulatory molecules and cytokines). Together, our experiments elucidate the differential immunogenicity of BT- and AKK- derived peptides and the orthogonal specificity of responder T cells in *in vivo* settings, likely reflecting baseline precursor frequencies of peptide-specific T cells.

**Fig. 4:**
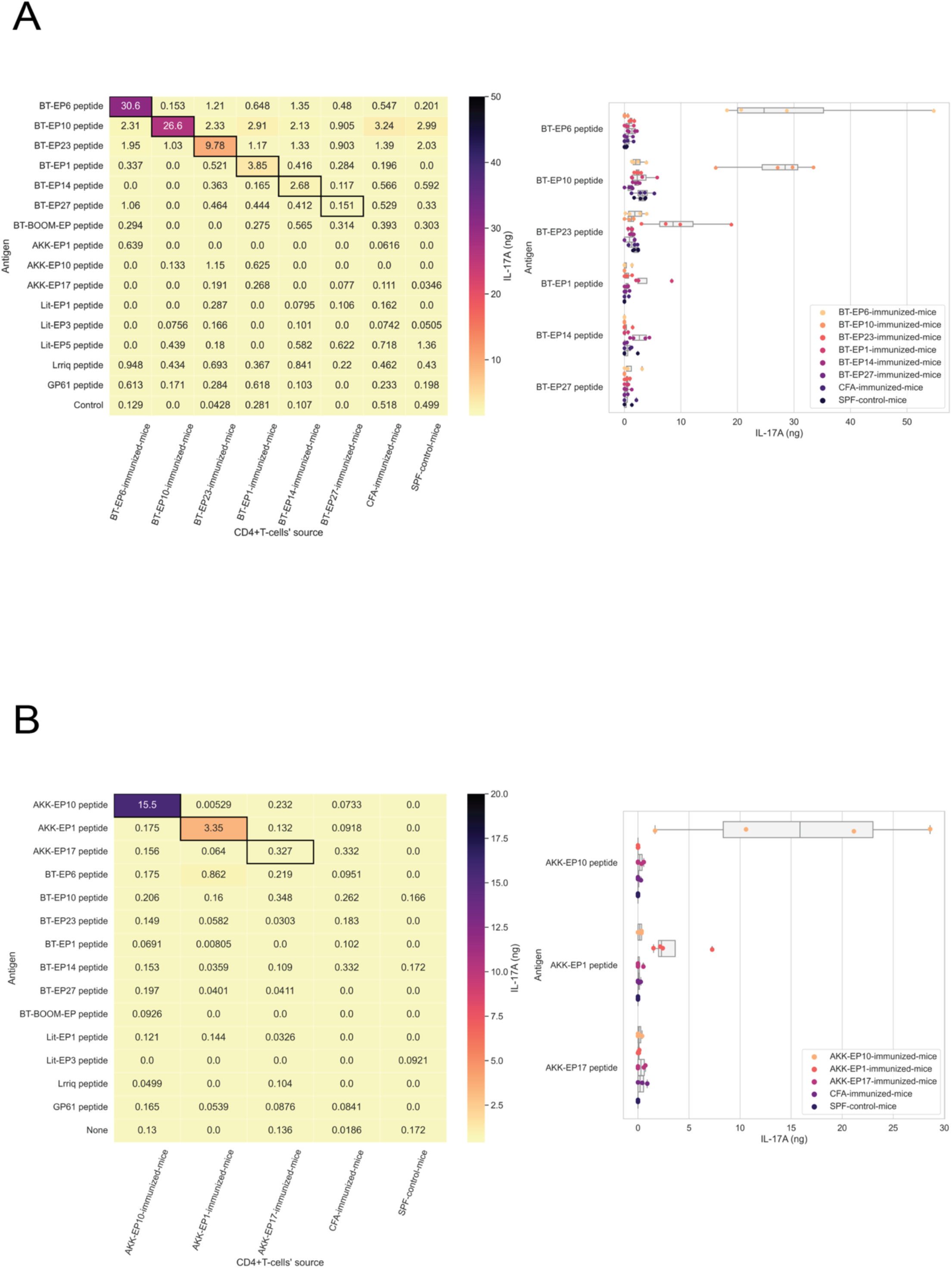
Orthogonal specificity and differential immunogenicity of BT and AKK derived peptides. IL-17A cytokine production quantification after ex-vivo restimulation of CD4+ T cells from mice immunized with (**a**) BT and (**b**) AKK peptides in CFA, respectively. Controls are mice immunized with CFA only and naïve nonimmunized mice. CD4+T cells from spleen and lymph nodes are sorted and restimulated with bone marrow derived dendritic cells (BMDCs) pulsed with an array of BT, AKK, mammalian, viral, and literature peptides, each individually. IL-17A ELISA is performed on co-culture supernatant after 3 days. Heatmap (left panel) depicts averaged IL-17A quantity in nanogram/ml (4 mice per peptide immunization). Boxplot (right panel) shows individual quantitative values. Data are representative of two independent experiments.

### Conservation of microbiota-derived peptides

Previous reports have proposed that T cell immune surveillance against the intestinal microbiota follows a model where T cell specificity is directed toward widely conserved epitopes shared by cohorts of colonists within a phylum. Particularly, it was shown that a T-cell hybridoma specific to a peptide derived from the BT protein BT0900 cross-reacted to two additional *Bacteroides* species, namely *B*. *ovatus* and *B*. *finegoldii* ^51^. Furthermore, a peptide from the β-hexosaminidase enzyme, conserved in members of the Bacteroidetes phylum, drove the differentiation of CD4+CD8αα+ intraepithelial lymphocytes (CD4IELs) ^53^. Additionally, commensals of the Firmicutes phylum were shown to harbour a conserved peptide of the substrate-binding protein (SBP) that induced T_H_17 and T_regs_ cells and was recognized by 13 different TCRs ^29^.

Thus, we sought to investigate the conservation of our epitopes across different bacterial strains and phyla. By conducting a BLAST-based epitope similarity analysis, our results revealed that BT-EP1 is highly conserved among the Bacteroides genus **(Fig. 5a)**. Specifically, BT-EP1 featured perfect homology in several strains such as Bacteroides ovatus, Bacteroides pyogenes, Bacteroides fragilis, and Bacteroides finegoldii, among many others. Additionally, BT-EP1 manifested amino acid sequence polymorphisms in Bacteroides stercoris, Bacteroides gallinaceum, and Bacteroides uniformis. Notably, BT-EP1 is also highly conserved among other families within the Bacteroidetes phylum and is found in the Verrucomicrobiota and Firmicutes phyla **(Fig. 5c,d)**.

**Fig. 5:**
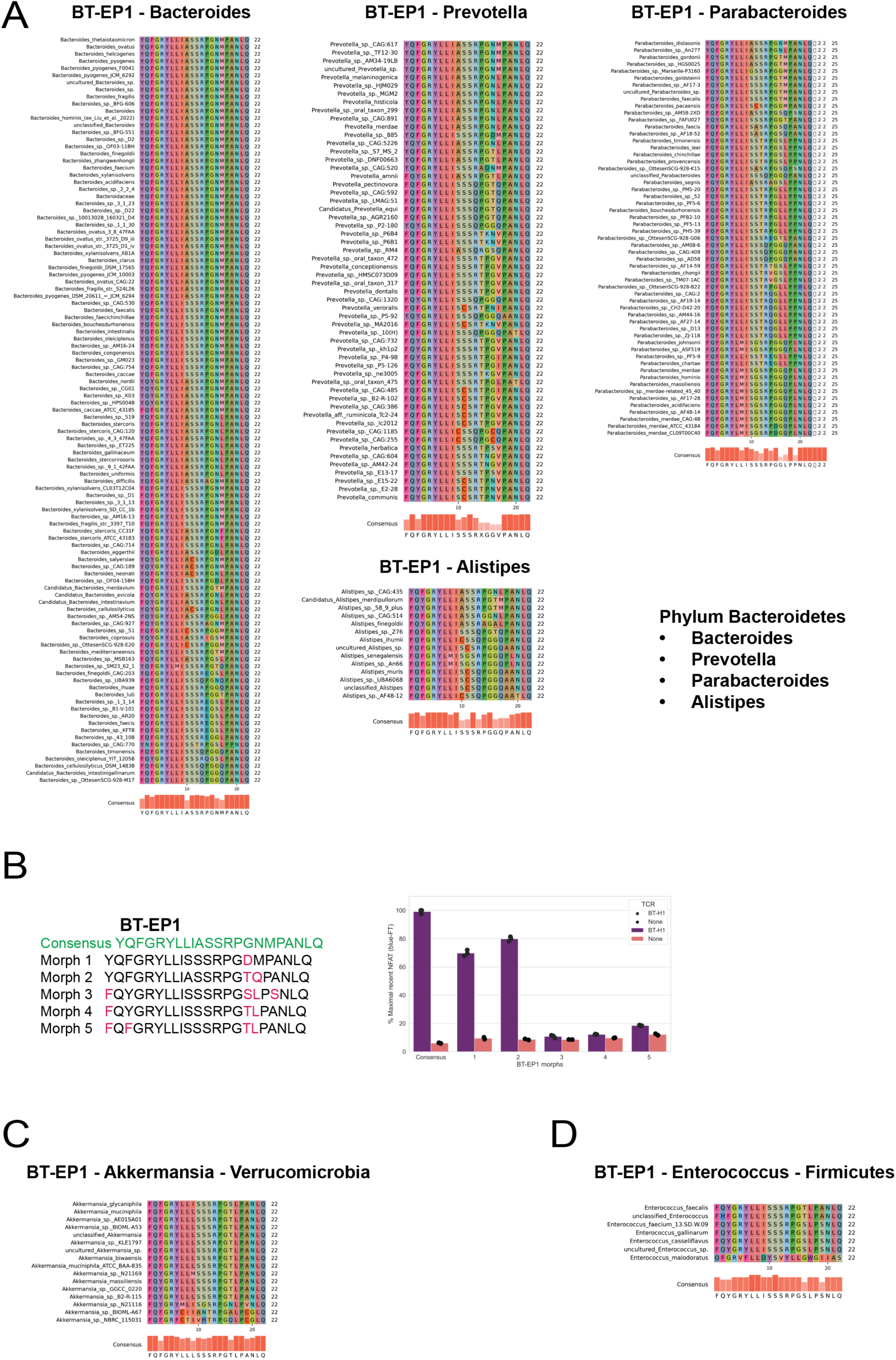
Conservation of microbiota-derived peptides. a, Multiple sequence alignment (MSA) of BT-EP1 in different families of the Bacteroidetes phylum of bacteria. **b,** Co-culture of BT-H1 with BT-EP1 polymorphs expressed in MCR reporter cells in 5:1 ratio. NFAT activation is measured after 16 h. BT-EP1 conservation in Verrucomicrobiota phylum (**c**) and Firmicutes phylum (**d**) of bacteria.

Subsequently, we examined whether the polymorphic forms of BT-EP1 were recognized similarly to the consensus BT-EP1 peptide. For this, we co-cultured BT-H1 with MCR+ reporter cells expressing various BT- EP1 polymorphs. Our findings indicated that certain polymorphs (specifically, numbers 3, 4, and 5) fully abolished the NFAT signal, while others (numbers 1 and 2) showed a reduction of 30% and 20% in the NFAT signal, respectively, compared to the consensus BT-EP1 peptide **(Fig. 5b)**. We then systematically probed the conservation of additional immunogenic peptides derived from BT and AKK bacteria. Our analyses showed that most peptides exhibit intra-phylum conservation, though the extent of epitope degeneracy varied **(Supplementary** Fig. 9b,d**,e)**. For instance, BT-BθOM-EP exhibited perfect homology in only five strains, whereas BT-EP23 showed perfect homology in 26 Bacteroides strains **(Supplementary** Fig. 9c,b). Comparatively, AKK peptides exhibited a lower degree of conservation when contrasted with BT peptides **(Supplementary** Fig. 9g,h). Together, these data suggest that epitope degeneracy is a common characteristic of microbiota-derived peptides, potentially selected for optimal T cell recognition and immune surveillance.

## Discussion

Mapping the intricate interactions between immune and microbial cells holds significant implications for deciphering immune regulation. T-cell responses at the antigen and epitope levels are highly complex, particularly given the body’s tremendous diversity of microbiota composition. Dissecting the microbial antigenic landscape is essential for interrogating the mechanisms underlying T cell immunodominance. Efforts aimed at comprehensively identifying antigenic epitopes have culminated in the establishment of the Immune Epitope Database (IEDB), an extensive repository of experimentally validated epitopes ^60,61^. However, despite notable advancements in gut microbiota biology, the number of validated microbial epitopes remains considerably lower compared to extensively studied organisms like Mycobacterium tuberculosis (MTB). For instance, the number of validated epitopes for MTB surpasses those derived from two of the most abundant and extensively studied gut microbes, AKK and BT, by over a hundredfold. The paucity of validated microbial epitopes potentially limits our full understanding of T cell and microbiota dynamics at mucosal surfaces. This underscores the need for studies that identify new antigens to enhance our insights into immune-microbiota interactions and their implications for human health.

In this study, we employed the MCR system to delve into the antigen-specific interactions between T cells and the gut microbiota, particularly focusing on AKK and BT bacteria. Our strategy to immunize mouse lungs with commensals confers several benefits. It facilitates the unbiased uptake of total antigenic material, enabling the processing and presentation of microbial epitopes by antigen-presenting cells. Simultaneously, it robustly activates and accumulates specific CD4^+^ T cells of functionally distinct immune phenotypes in high numbers. By constructing T cell hybridomas derived from these activated/reactive T cells and subsequently screening against comprehensive, unbiased, and genome-wide MCR libraries of AKK and BT bacteria, we aimed to uncover the identities of *in vivo* presented microbial peptides. We showed that bacteria-reactive T cell hybridomas exhibit polyreactivity towards select epitopes at the TCR-MCR interface, as evidenced by induction of NFAT and Nur77 signaling in MCR^+^ cells. Our endeavors resulted in the identification of 27 novel antigenic epitopes—13 from AKK and 14 from BT—thus expanding the repertoire of recognized bacterial antigens. We further demonstrated that T cell hybridomas specific for viral and mammalian species recognize our identified BT and AKK-derived peptides, suggesting that these MCR-fused peptides have a greater capacity to bind a wide range of TCRs of varied affinities and specificities. We observed that neither the MCR-fused bacteria peptides nor synthetic peptides activated the bacteria-reactive T-cell hybridomas. However, performing *in vivo* mouse immunizations with these peptides derived from AKK and BT bacteria showed their immunogenicity, as evidenced by the secretion of IL- 17A and IFN-γ cytokines in recall assays. Notably, *ex vivo* stimulated T cells specific for one peptide did not respond to other peptides, differing from the results observed in the MCR co-cultures. Of note, we demonstrated the conservation of these peptides across a wide array of bacterial strains within the same bacterial phylum or even across different phyla.

TCR interaction with peptides presented by MHC molecules (pMHC) is the modus operandi of T cell activation and differentiation and constitutes the primary basis for specific immunity to microbiota-derived antigens. T cell cross-reactivity is a well-documented phenomenon in the immune system, characterized by the ability of a single T cell to recognize multiple pMHCs via the TCR ^62^. This degenerate recognition is viewed as an intrinsic property of the immune system, essential for a robust immune response. However, TCR cross-reactivity can also precipitate autoimmune diseases through the recognition of self-peptides ^63^. In our *in vitro* MCR co-cultures, we observed widespread polyreactivity of AKK- and BT-reactive T cell hybridomas with select AKK- and BT-derived epitopes. However, these observations were not mirrored in *in vivo* immunization experiments. We assume that these contradictory results may stem from the nature of T cell activation itself. TCR-pMHC binding is the first step in the T cell activation cascade, requiring the ligation of co-stimulatory molecules and the provision of cytokines as the second and third signals, respectively, for full activation. Consequently, we argued that such polyreactive T cells do not receive the second and third signals provided *in vivo*. Another possibility could be that such polyreactive T cells are rare in nature and their proliferation is tightly regulated *in vivo.* Of note, cross-reactivity of CD4+ T cells toward certain species of bacteria and yeast has been documented ^16,57,64^. Further research is needed to understand the mechanisms controlling the activity of cross/poly-reactive and their potential implications for immune responses against the microbiota.

Having observed that the synthetic peptides do not activate the bacteria-reactive T cell hybridomas, we first examined the possibility of weak binding of the peptides to the MHC. Even when supra-physiological amounts of the peptides were used, these observations of non-binding persisted. Furthermore, the TCR signaling machinery was intact in the T cell hybridomas, excluding any confounding factors stemming from defective TCR signaling cascades. The only exception was the bidirectional interaction between the BθOM TCR_H_ and BθOM MCR or BθOM synthetic peptide. Moreover, these peptides do not act as antagonists for TCR signaling. Our findings suggest that the MCR-fused peptides are recognized in a novel structural conformation, which does not translate into effective T-cell hybridoma activation. Furthermore, these peptides have a binding mode which is not resembled *in vivo*.

Notably, the phenomenon of TCR-MHC biophysical binding without subsequent T cell signaling has been reported in literature ^65^. Specifically, it was shown that a single TCR could bind peptides in different binding topologies of the TCR-peptide-MHC complex, with some peptides inducing TCR signaling and others not. The latter were revealed to have a comparable affinity to TCR agonists but did not instruct an activating or inhibitory signal, possibly due to their unconventional orientation, which is incompatible with T-cell signaling. Our MCR- fused peptides manifest similar patterns of TCR binding with no stimulation or inhibition of TCR signaling, likely due to their unconventional pMCR structure. These findings may point to a systematic caveat in our experiments. Our strategy to generate comprehensive bacterial peptide libraries, as opposed to a biased and restricted set of proteins, might have enriched for such peptides. Further investigation is required to understand the common structural denominator of these peptides and the lack of signaling. Understanding the biological implications of these observations may reveal new aspects of immune regulation and response to gut bacteria.

Numerous bacteria-derived superantigens have been characterized in relation to T cells ^66–70^. These superantigens induce a hyper-stimulatory response by cross-linking TCR Vβ with MHC class II α1 molecules on antigen-presenting cells (APCs). This interaction leads to vigorous T cell proliferation and cytokine production. Additionally, B cell superantigens expressed by commensal bacteria in the human gut microbiota have been reported to stimulate IgA production and bacterial coating ^71^. Our identified peptides, presented by MCR structures, exhibit characteristics reminiscent of superantigens. Specifically, these peptides bind a diverse range of TCRs with bacterial, viral, and mammalian reactivities and affinities, in an MHC-restricted manner. For this reason, we refer to these peptides as “super-peptides. In contrast to classical superantigens, which bind and induce signaling in T cells, our super-peptides bind to T cells without triggering a signaling response. This represents a significant distinction between our super-peptides and traditional superantigens. Notably, we have previously reported a few examples of super-peptides derived from mammalian sources ^1^. We hypothesize that the MCR forms a structural configuration that does not resemble the conventional MHC-peptide complex, suggesting that these peptides probably don’t represent the nominal T cell specificity. Furthermore, we speculate that the MCR may stabilize these peptides in their full format, which contrasts with *in vivo* conditions where peptides undergo antigen processing and presentation. Overall, our findings offer an intriguing example of potential T cell super- peptides derived from gut bacteria. Future research is needed to explore the physiological relevance of these super- peptides.

Despite the above-mentioned complications, our immunization experiments showed that the bacteria-derived epitopes identified in our screen can be immunogenic, mounting both a Th1 and Th17 response. IL-17A plays a vital role in maintaining the integrity and function of mucosal surfaces, particularly in the gut. As a cornerstone cytokine, IL-17A is essential for the host’s defense against pathogens and for maintaining a balanced relationship with the gut microbiota. It enhances epithelial barrier function by inducing the production of antimicrobial peptides, promoting mucus secretion, and strengthening tight junctions between epithelial cells, thereby supporting a robust mucosal barrier. However, its dysregulation, influenced by gut microbiota imbalances, can lead to significant immune pathologies, highlighting the necessity for precise regulatory mechanisms ^72^. On the other hand, studies have shown that IFN-γ helps balance immune responses at mucosal surfaces, ensuring that commensal microbes are tolerated while pathogenic microbes are effectively controlled ^73^. However, its overproduction in response to microbiota-derived peptides can contribute to chronic inflammation and diseases such as inflammatory bowel disease (IBD) ^74^. Our findings that T cells responding to the identified epitopes produce both IL-17A and IFN-γ underscore the importance of the balance between these cytokines. Further investigation is warranted to understand how their dysregulation contributes to immune-mediated diseases.

Poised with trillions of microbial species and an ocean of antigens, the immune system must concoct ingenious and sophisticated strategies to navigate this complexity while managing the energy demands required for species-specific recognition. Our data, in alignment with prior research, suggest that the immune system is prudent in its recognition of bacterial species. It may prioritize the identification of highly conserved antigens shared among evolutionarily linked bacterial species, thereby minimizing the energy expenditure associated with recognizing individual bacterial strains. This strategic allocation of resources may enable the immune system to efficiently cope with the microbiota while conserving energy to target and eradicate pathogens.

In conclusion, this study provided novel insights onto bacteria-derived peptides and addressed a critical knowledge gap on identifying immunogenic bacterial epitopes. Our results may help in understanding the reactivity of T cell responses to gut bacteria and form the basis for future antigen screens.

## Limitations of our work

In our endeavor to explore *in vivo*-educated T cell specificities, we generated a limited number of AKK- and BT- reactive T cell hybridomas, representing only a fraction of the T cell repertoire. Nevertheless, we were able to identify peptide ligands for these T cell clones, which showed intriguing polyreactive and conserved-epitope reactive features. However, it would be very dangerous to extrapolate from this low number of clones to broad statements about microbiota-reactive T cells. Future efforts could leverage tetramer technology in conjunction with single-cell RNA sequencing (scRNA-seq) and single-cell TCR sequencing (scTCR-seq) to achieve finer resolution in dissecting the phenotype and total specificity of T cell responses. Second, we focused solely on IL- 17A and IFN-γ and did not analyze other cytokines such as IL-21, IL-22, IL-10, and IL-4. Third, our study centered on two strains from two bacterial phyla, and it remains unclear whether what we observe is typical of immune responses against other bacterial phyla. Fourth, we could not explore the functional relevance of the TCR- peptide interactions that we observed. For this, we would require TCR-transgenic systems that do not yet exist. Therefore, the relative contribution of broadly-conserved-peptide-reactive T cells to mucosal immunity remains unclear. Finally, we could not address the relevance and dynamics of peptide reactive T cells in inflammatory and autoimmune disease conditions in the timeframe of this study. Exploring whether similar findings are true in humans, and whether such broadly conserved epitopes play a role in T-cell -mediated inflammatory bowel diseases is a major future area of interest. Addressing the aforementioned limitations should enhance our understanding of mucosal immune responses.

## Materials and methods

### Mice

C57BL/6J SPF mice were reared at the ETH Zürich mouse facility (EPIC), Switzerland. C57BL/6 GF mice were raised in surgical isolators under strict hygiene conditions at the ETH Phenomic Center. They were routinely tested for contamination using aerobic and anaerobic cultivation, culture-independent methods to assess intestinal bacterial densities, and serology/PCR tests for common viruses and eukaryotic pathogens. Experiments were performed with sex and age matched animals, ranging from 6-15 weeks. Mice were euthanized with CO_2_ or pentobarbital. All animal experiments were performed according to the institutional guidelines and Swiss federal regulations and were approved by the animal ethics committee of Kantonales Veterinärsamt, Zürich, Switzerland (permission no. ZH104/21, ZH134/18, ZH074/23.).

### Immunization with bacteria

Bacteroides thetaiotaomicron VPI-5482 and Akkermansia muciniphila YL44 strains were grown anaerobically (5% H_2_, 10% CO_2_, rest N_2_) at 37°C, overnight in brain heart infusion (BHI) supplemented media (BHIS: 37 g/L BHI [Sigma]; 1 g/L-cysteine [Sigma]; 1 mg/L Hemin [Sigma]). Bacteria were centrifuged and pellets were kept frozen at -20 until used. Heat killing (HK) was performed by incubating bacteria for 30 minutes at 60°C. Mice were immunized twice intratracheally (i.t.) with HK bacteria, with a dose of 10^7 colony forming units (CFUs) per mouse.

### Immunization with peptides in CFA/IFA

Peptides were purchased with >95% purity from Genosphère Biotechnologie (92100 Boulogne-Billancourt, France). 100 μg of peptide in PBS was mixed 1:1 with Complete Freund’s Adjuvant (CFA - Difco Laboratories) and was injected subcutaneously (s.c) into SPF mice on one flank. Mice were challenged with 50 μg of peptide mixed 1:1 with Incomplete Freund’s Adjuvant (IFA - Difco Laboratories) injected s.c. on the other flank.

### Preparation of MCR libraries

Bacterial genomic DNA (gDNA) was sheared using Takara fragmentation kit (Takara Bio) and run on 2% agarose gel. Fragments of an average of 150 bp were excised and purified using Zymoclean Gel DNA Recovery Kit (Zymo Research). Purified DNA was ligated with 100 bp linkers and ligation product was run on 2% gel at 70 volts. The smear above 250 bp was excised from the gel, purified and PCR amplified (20 cycles using Phusion High–Fidelity DNA Polymerase). PCR product was purified and cloned into the pMY-MCR2-I-A^b^ plasmid using Gibson assembly, and product was transfected into One Shot™ TOP10 Electrocomp™ E. coli (Thermo Fisher Scientific), generating 2.5 × 10^7^ clones for BT bacteria and 1.17 × 10^7^ clones for AKK bacteria.

### Retroviral transfection and transduction of reporter cells

Phoenix packaging cell line was used for production of retroviruses. Cells were cultured in IMDM complete containing 10 %FCS, 1% P/S and 50 μM b-Mercapto-EtOH and were 70-80% confluent at time of transfection. At day 3, medium containing virus was harvested, centrifuged at 10.000 g for 1 hour at 4°C. Concentrated virus was used for spin infection of 16.2X MCR reporter cells in complete medium containing Polybrene. 16.2X cells were FACs analysed after 2-3 days for expression of CD3e and/or MHC II. MCR+ cells were sorted on BD FACSAria Fusion - BSL2 sorter.

### Sorting of activated T cells and generation of hybridomas

Immunized or naïve mice were perfused with PBS prior to lung harvest. Lung organ was dissociated using gentle MACS (Miltenyi Biotec) and incubated with 600 U/mL Collagenase IV (Worthington) and 200 U/mL DNase I (Sigma-Aldrich) for 45 min at 37°C. Tissue was homogenized again with gentle MACS and passed through a 40 μm cell strainer. RBCs were eliminated by ACK and single-cell suspension was filtered through a 70 μm cell strainer. CD44 high CD62L low CD4+T cells were sorted on a FACSAria Fusion - BSL2 sorter. Subsequently, T cells were activated using plate bound aCD3 antibody (2 μg /ml) and supplemented with anti-CD28 (2 μg /ml) and mouse IL-2 (20 ng/ml) for 36 hours. Activated T cells were then fused with the TCRα-β- BW5147 fusion partner in a 1:1 ratio using polyethylene glycol (PEG)1500 (Thermo Scientific Chemicals). Fused cells were cultured in selection medium containing HAT supplement (1/50 dilution – purchased from Gibco) and ciprofloxacin antibiotic for 10 days. Growing clones were checked for TCRb expression.

### T cell–MCR co-culture

CD4+T cell hybridomas or primary CD4+T cells were co-cultured with MCR+ reporter cell line in a 5:1 ratio. Recent NFAT activation was measured after 16 h of co-culture.

### Generation of bone marrow derived dendritic cells (BMDCs)

Bone marrow cells harvested from naive SPF C57B/6J mice were *in vitro* differentiated with granulocyte-macrophage-colony-stimulating factor (GM-CSF) for 7-9 days.

### ELISA

Primary CD4+T cells were co-cultured with BMDCs pulsed with 10 μg/ml of peptide in a 5:1 ratio. Culture supernatant containing cytokines was harvested after 36 hours and subjected to IL-17A and IFN-γ ELISAs. Coating of flat bottom ELISA plates (NUNC) was performed with 5μg/ml of antibody in PBS with the following antibodies: IFN-γ monoclonal antibody, clone R4-6A2 (catalog # 14-7312-85, eBioscience™) and anti-mouse IL- 17A antibody, cloneTC11-18H10.1 (catalog # 506901, BioLegend®). For standards, 50 ng/ml of the following proteins were used: mouse IFN-γ recombinant protein (catalog # 14-8311-63) and mouse IL-17A recombinant protein (catalog # 14-8171-62, eBioscience™). Biotinylated antibodies were biotin IFN-γ monoclonal antibody, clone XMG1.2 (catalog # 13-7311-85, eBioscience™) and biotin anti-mouse IL-17A antibody, clone TC11-8H4, (catalog # 507001, BioLegend®). 4-Nitrophenyl phosphate disodium salt hexahydrate (pNPP - Sigma-Aldrich) tablets solved in Diethanolamine buffer was used as substrate for Streptavidin-AP (catalog # 7105-04, SouthernBiotech).

### Amplicon NGS library preparation and data analysis

RNA was extracted from 500k MCR reactive reporter cells from specified enrichment rounds and converted to complementary DNA (cDNA). Two-step PCR approach was conducted for library amplification using standard protocol. The primers included locus-specific sequence, i.e., peptide containing fragment in the MCR vector and Illumina universal adaptors. PCR product was purified using Zymo DNA-clean and concentrator. Illumina MiSeq machine was used for sequencing (Microsynth AG- Switzerland). FastQC (v0.12.1) was used for RNA sequencing data quality assessment ^75^. Bbmap, Flash & Pandaseq softwares ^76–78^ were used for merging overlapping paired-end reads and sequence of peptide harboring fragment was subsequently obtained. The following criteria were used to ensure high quality data: sequences must contain a DNA-encoding peptide sequence and length of peptide is dividable by 3 and leaves no remainder. Normalization was done using size factors- i.e., number of reads per sequence divided by total number of reads per sample.

### Data and code availability

Raw NGS sequencing files will be deposited in the NCBI Gene Expression Omnibus and will be publicly available upon publication. The following Python packages were used for data analysis: pandas, matplotlib, and seaborn. The lead contact can provide any additional information needed to reanalyze the data reported in this paper upon request.

### Multiple sequence alignment (MSA) analysis

Clustal analysis was done at https://www.ebi.ac.uk/jdispatcher/msa/clustalo. MSA was visualized using pyMSAviz Python package.

### Surface and intracellular staining and flow cytometry

Single cell suspensions were stained with Fixable Viability Dye eFluor 780 (eBioscience, 65-0865-18) to exclude dead cells and Fc block (anti-CD16/CD32; 2.4G2; home-made) was used to prevent non-specific antibody binding, in both surface and intracellular staining. The following antibodies were used: CD3e (catalog # 100348, BioLegend), CD4 (catalog # 100447, BioLegend), CD44 (catalog #103044, BioLegend), CD62L (catalog # 741924, BD Biosciences), CD11c (catalog # 565872, BD Biosciences), IL-17A (catalog # 506912, BioLegend), IFN-γ (catalog # 557649, BD Biosciences), TCRb (catalog # 109230, BioLegend), Nur77 (catalog # 12-5965-82, Invitrogen). Fixation and permealization was done using the eBioscience Foxp3/Transcription Factor Staining buffer set (ThermoFisher Scientific, catalogue no. 00-5523-00). Cells were acquired on BD FACSymphony A5 SE and Cytek Aurora and analyzed in FlowJo (BD Biosciences).

## Acknowledgements

We thank Emma C. Slack (ETH Zürich, Switzerland) for providing GF mice and bacterial strains and members of the Slack lab for coordinating experiments. We are grateful to the team of the ETH flow cytometry core facility for help with cell sorting and to the animal caretakers of the ETH Phenomics Center (EPIC) for mouse maintenance. We thank Drs. Michael Dustin and Andrew Macpherson for the discussion.

## Author contributions

A.S. conceived ideas, designed and performed experiments, analyzed data, and wrote the manuscript. E.R. helped A.S. set up the MCR system. D.H. and E.S. provided GF mice and bacterial strains. A.S., F.M., and M.K. discussed and interpreted data. M.K. conceived and supervised the project, reviewed and revised manuscript, and secured funding. All authors reviewed and approved the manuscript.

## Conflict of interest

The authors declare no competing financial interests.

---

**Supplementary Fig. 1:**
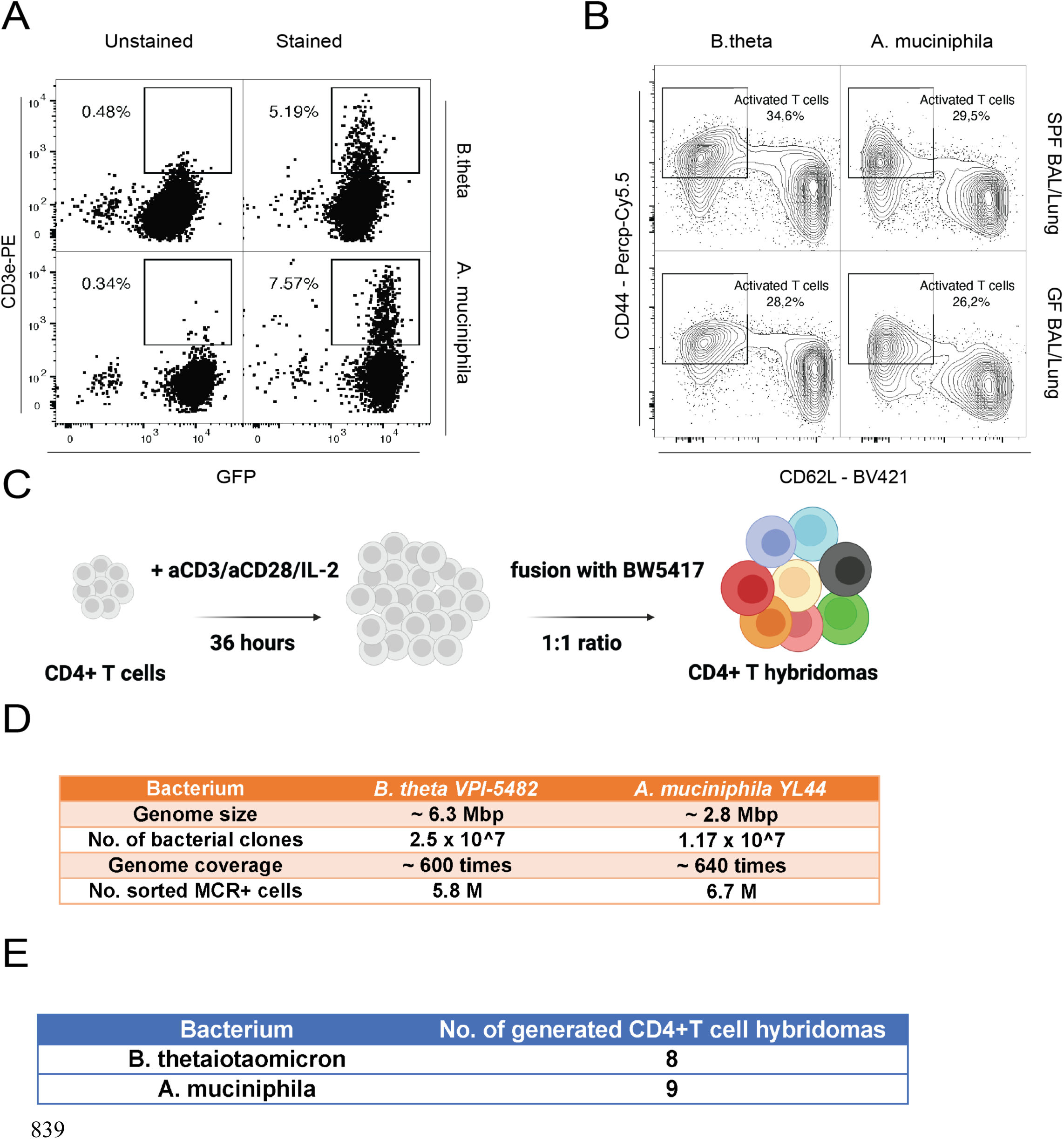
Coverage of MCR libraries and numbers of CD4+T cell hybridomas. a, Transduction efficiency of BT and AKK MCR libraries. MCR negative 16.2X reporter cells are transduced with MCR plasmids of BT and AKK bacteria. Depicted is percentage of MCR+ cells reporter cells. **b,** Sorting strategy for CD44hi CD62L low activated CD4+ T cells from immunized SPF and GF mice. **c,** Process for generation of CD4+T cells hybridomas. **d,** Summary statistics of bacterial genome coverage and number of sorted MCR+ cells in each library. **e,** Numbers of generated T cell hybridomas.

**Supplementary Fig. 2:**
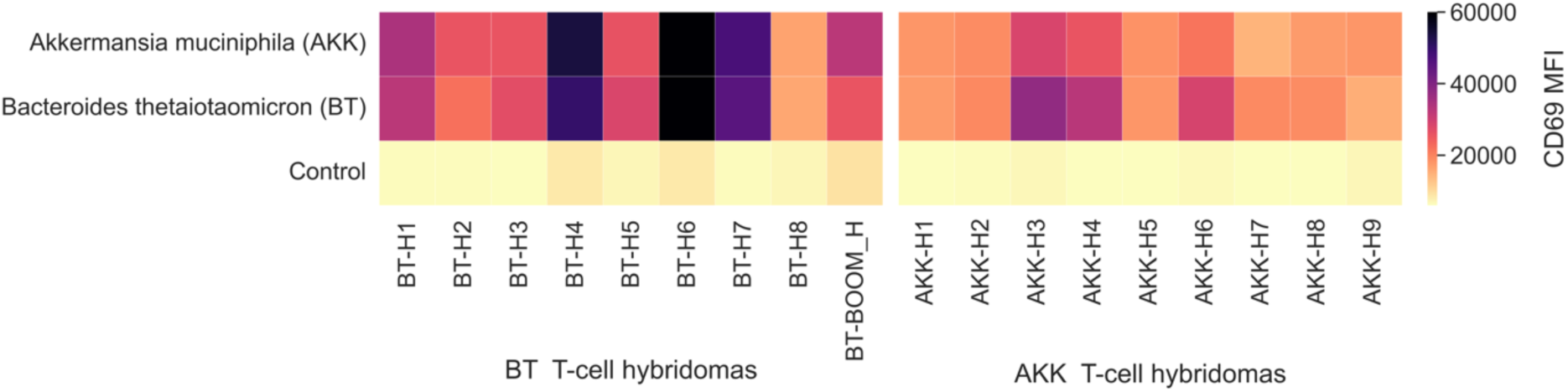
Reactivity of T-cell hybridomas toward different bacteria. Heatmap depicting reactivity of all generated T-cell hybridomas. T cell hybridomas are co-cultured with BMDCs pulsed with or without bacteria. CD69, an early marker of T cell activation, was measured after 24h. Colorbar represents MFI of CD69 expressed in T cells. The data are representative of two independent experiments.

**Supplementary Fig. 3:**
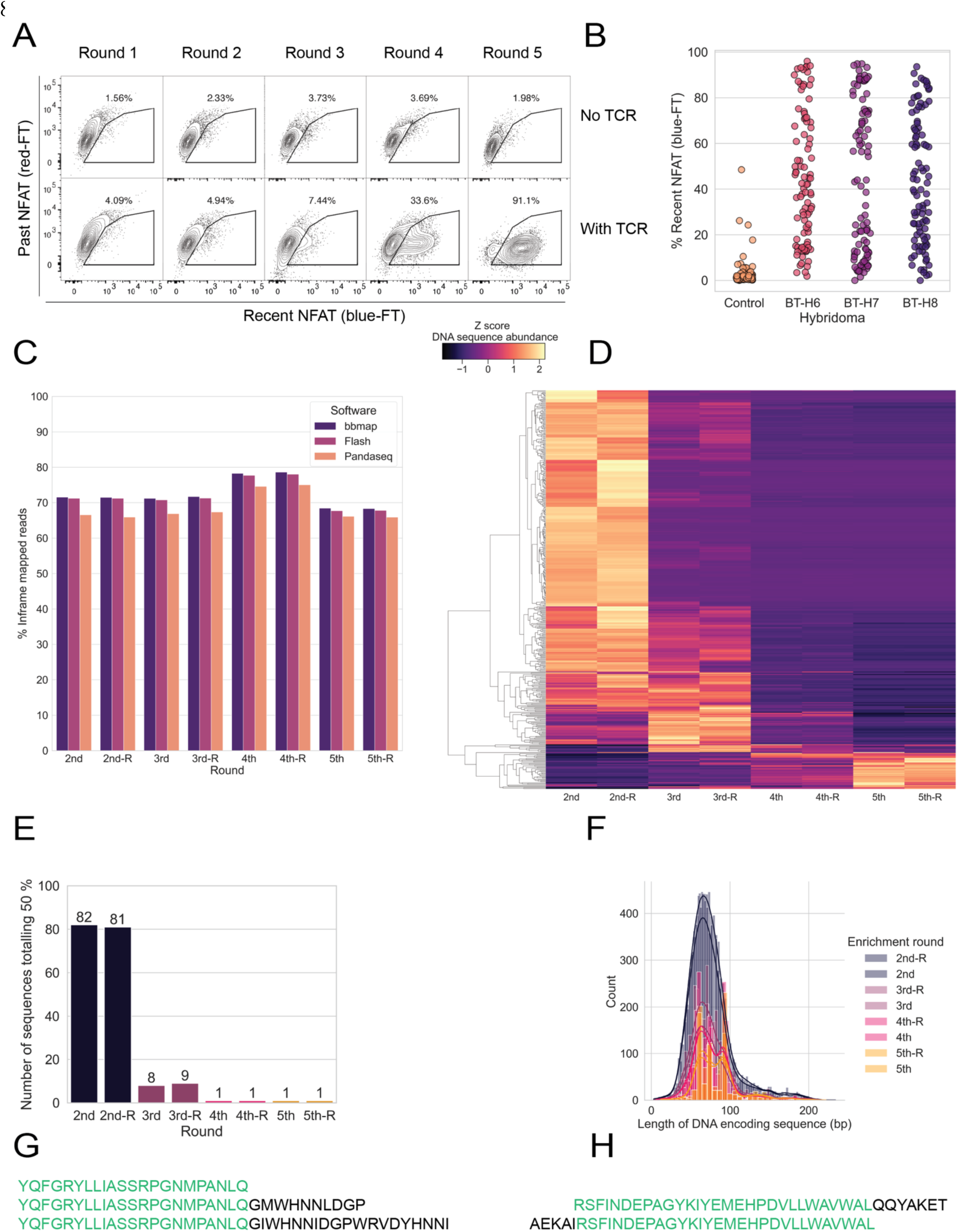
Representative enrichment of T cell hybridoma specific MCR+ reactive reporter cells. a, Iterative enrichment of target reporter cells. MCR library is left untreated or co-cultured with T cell hybridomas in a 1:5 ratio. Activated reporter cells are sorted, expanded for one week, and co-cultured again with the same T cell hybridomas until significant NFAT signal above background is reached. Number inside gate represents percentage of activated reporter cells. Data are representative of one independent experiment. **b,** Recall of MCR reactive reporter cells. Single reporter cells from the last round of enrichment are sorted, expanded and then co-cultured with T cell hybridomas initially used for enrichment. **c,** Alignment and mapping of NGS amplicons. Shown is the percentage of aligned DNA sequences that encode inframe peptides. Bbamp, Flash, and Pandaseq are three different softwares used to align NGS fastq files. **d,** Heatmap depicting NGS data of representative enrichment. Each row is a DNA-sequence encoding peptide. Colorbar represents Z score of row’s abundance (i.e., read counts). **e,** Number of DNA-sequence encoding peptides totaling 50 percent fraction of total read counts in each enrichment round. **f,** Overlayed length distribution of all DNA sequences in different enrichment rounds. **(g,h)** depict examples of two highly abundant encoded peptides found in different overlapping formats.

**Supplementary Fig. 4:**
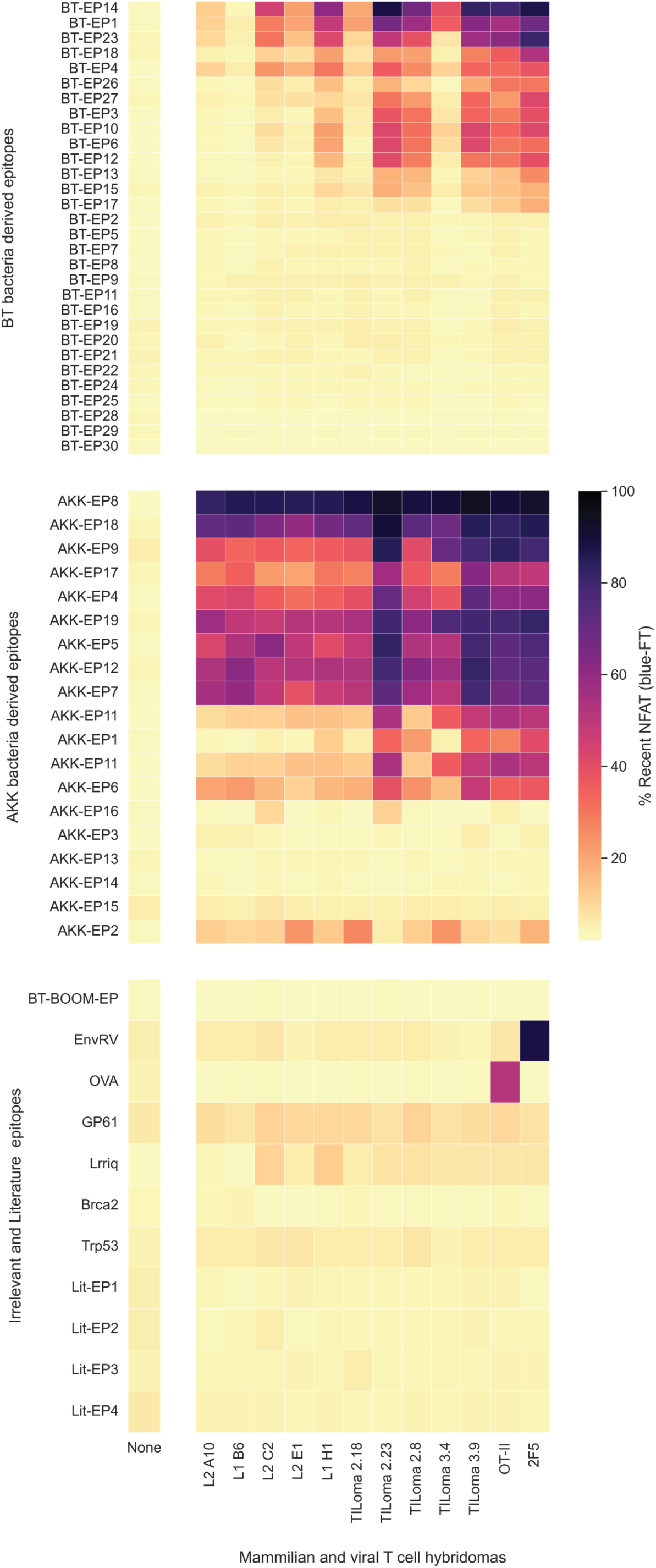
Mammalian and viral T cell hybridomas show polyreactivity to microbiota-derived epitopes in MCR co-cultures. Co-culture of T cell hybridomas of different sources with BT, AKK and irrelevant epitopes. T cell hybridomas (L2 A10, L1 B6, L2 C2, L2 E1, and L1 H1) are specific to influenza virus. “TILoma” denotes T cell hybridomas specific to the mammalian cancer cell line EO771. OT-II and 2F5 T cell hybridomas are specific to OVA and envRV epitopes, respectively. Each T cell hybridoma is co-cultured with every epitope individually and NFAT activation was measured after 16 hours. Colorbar represents percentage of recent NFAT activation. The data are representative of three independent experiments.

**Supplementary Fig. 5:**
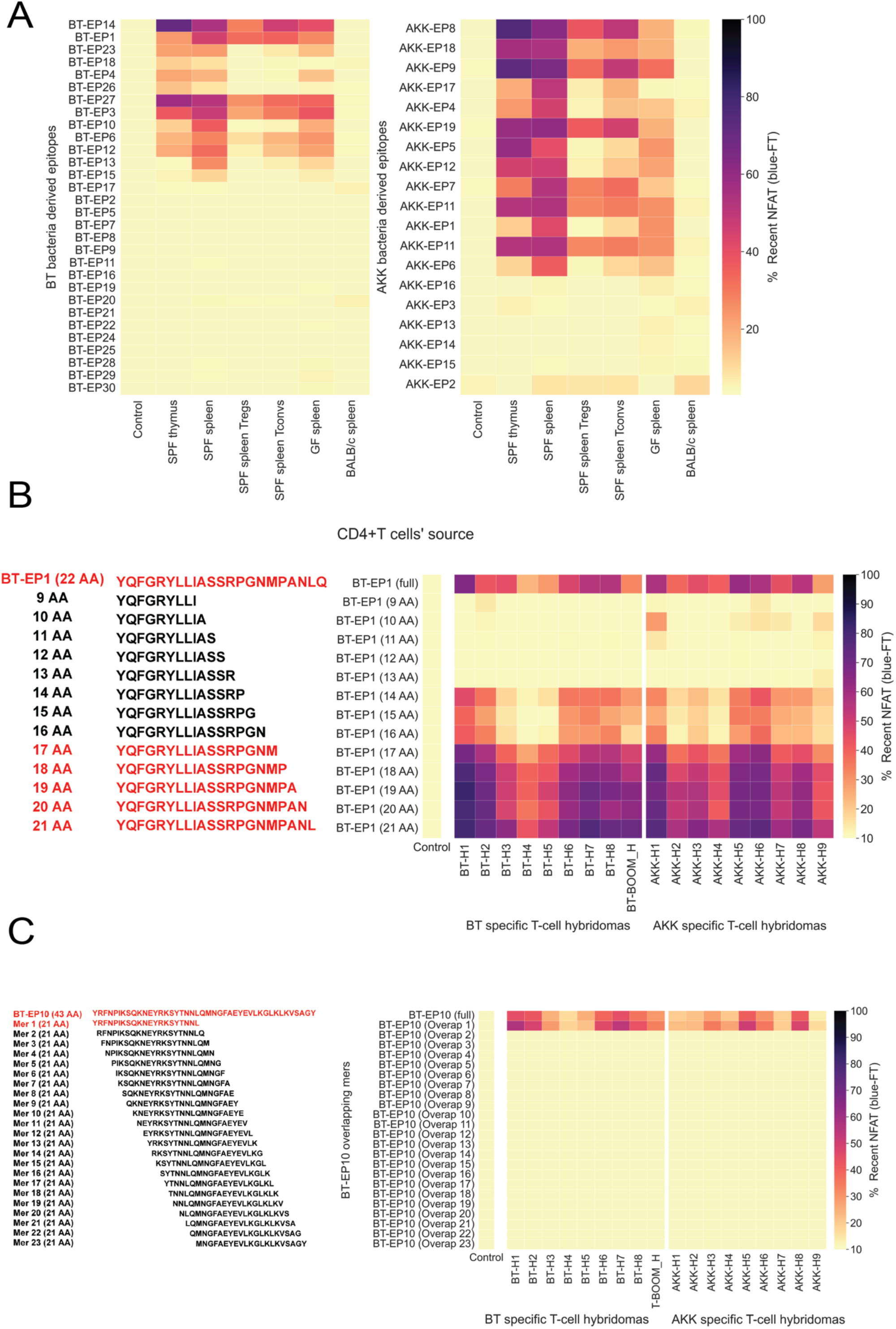
Steady state T cells recognize microbiota-derived peptides and identification of peptide core binding domain. a, Primary CD4+T cells from SPF, GF and Balb/c mice are co-cultured with BT- and AKK-derived epitopes in a 5:1 ratio for 16 h. Colorbar represents percentage of recent NFAT activation. **b,** Extending peptide fragments of BT-EP1 are co-cultured with BT- and AKK-reactive T cell hybridomas. Colorbar represents percentage of recent NFAT activation. **c,** Overlapping peptide fragments of BT-EP10 are co-cultured with BT- and AKK-reactive T cell hybridomas. Colorbar represents percentage of recent NFAT activation.

**Supplementary Fig. 6:**
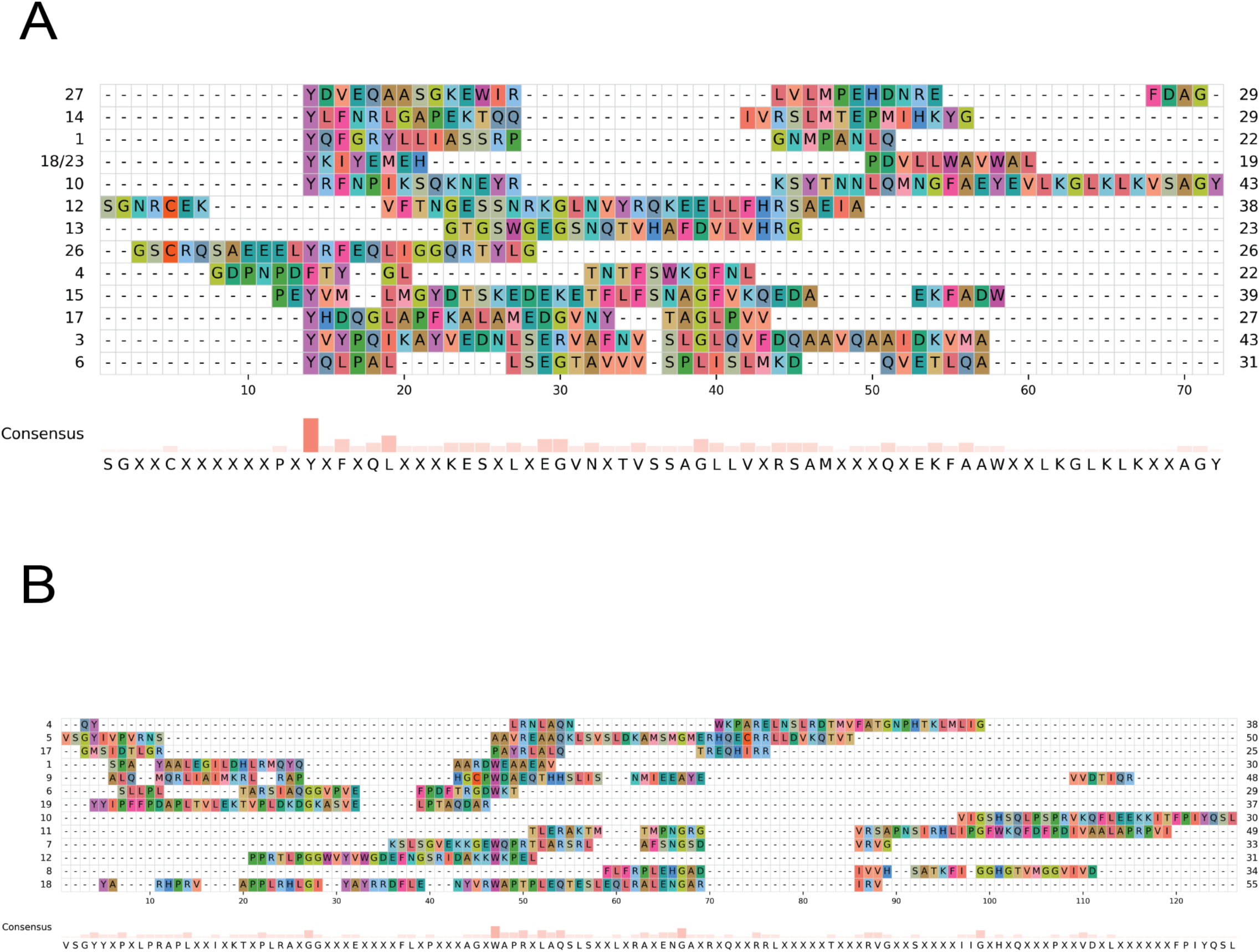
Clustal analysis of reactive BT- and AKK-derived peptides. Clustal analysis and visualization of reactive (a) BT-derived peptides and (b) AKK-derived peptides. Notice conservation of tyrosine (A) amino acid as position one.

**Supplementary Fig. 7:**
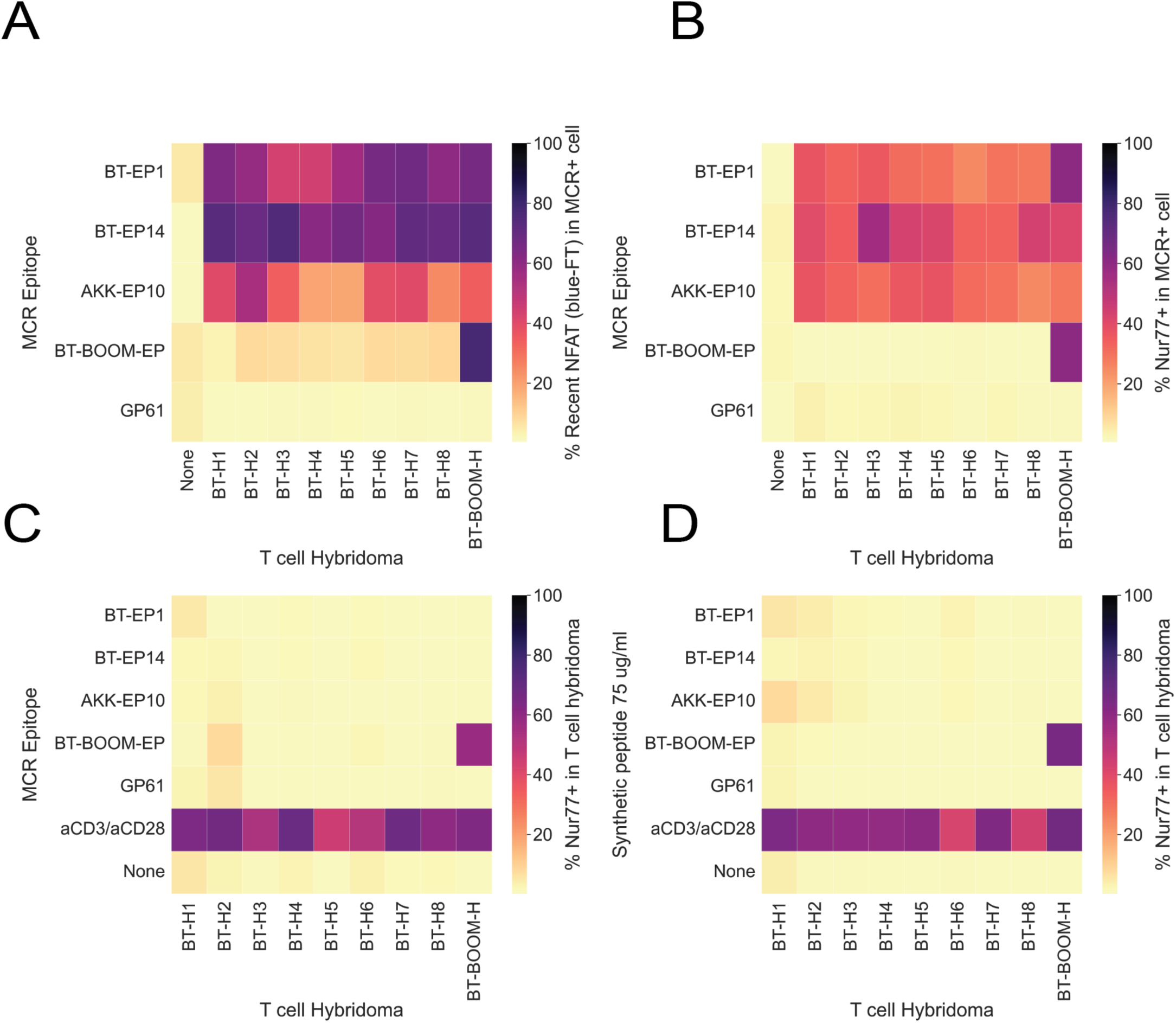
Microbiota-reactive T cell hybridomas are not activated by identified cognate peptides. Shown is co-culture of BT specific T cell hybridomas with BT, AKK, and irrelevant epitopes expressed in MCR cells or pulsed in BMDCs. Each T cell hybridoma is co-cultured with each epitope individually. **a,** Co-culture of T cell hybridomas with epitopes expressed in MCR cells. NFAT activation is measured after 16 hours. Colorbar represents percentage of recent NFAT activation in MCR+ cells. **b,** Co-culture of T cell hybridomas with epitopes expressed in MCR cells. Nur77+ upregulation is measured after 4 hours. Colorbar represents percentage of Nur77 in MCR+ cells. **c,** Co-culture of T cell hybridomas with epitopes expressed in MCR cells. Nur77+ upregulation is measured after 4 hours. Colorbar represents percentage of Nur77 in T cell hybridomas. **d,** Co-culture of T cell hybridomas with synthetic epitopes pulsed in BMDCs. Nur77+ upregulation is measured after 4 hours. Colorbar represents percentage of Nur77 in T cell hybridomas. The data are representative of two independent experiments.

**Supplementary Fig. 8:**
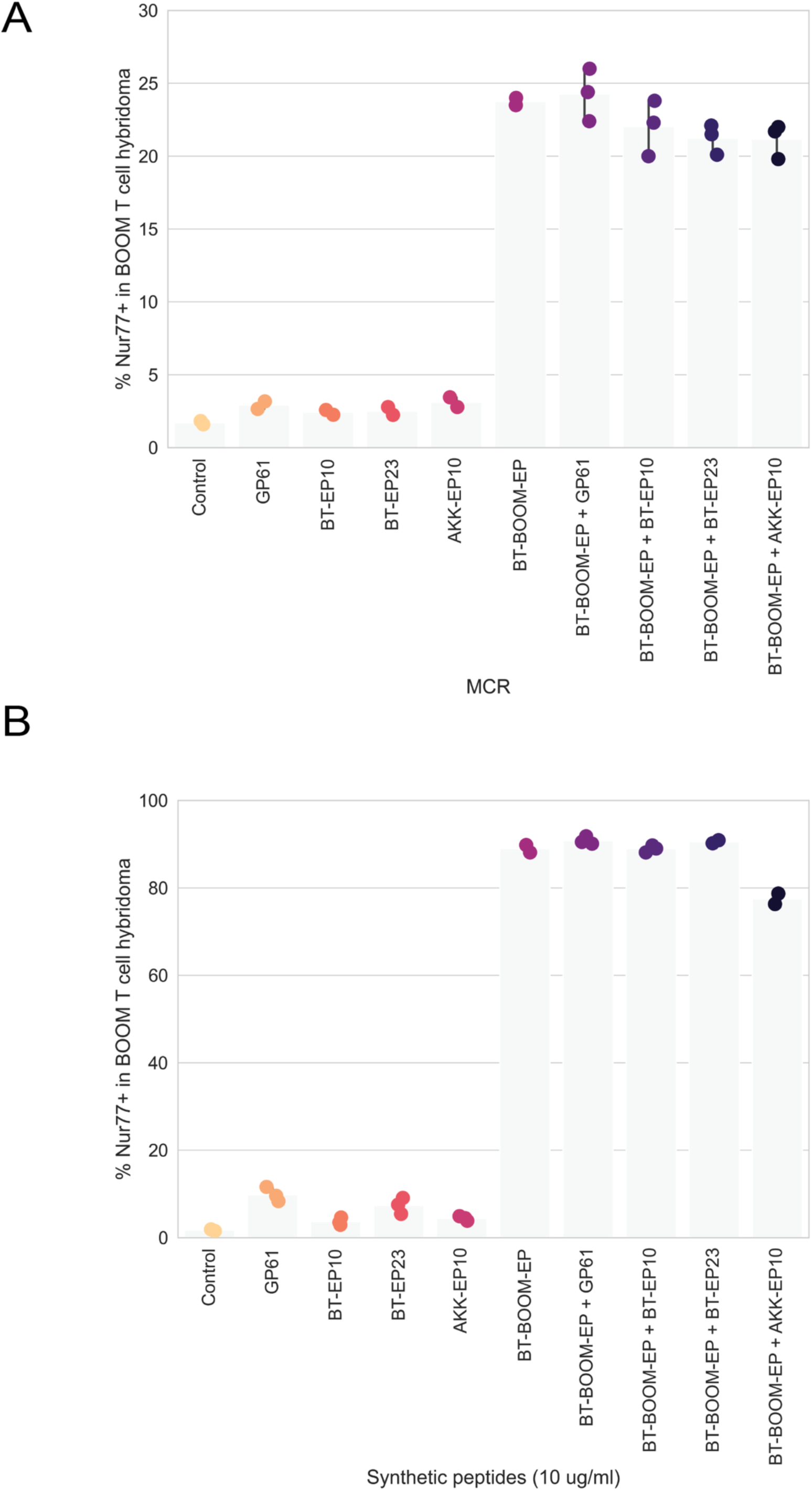
Microbiota-derived epitopes do not antagonize cognate T cell hybridomas. Co-culture of BθOM T cell hybridoma with an array of epitopes in different configurations, either expressed in MCR system (**a**) or pulsed in BMDCs (**b**). Nur77+ upregulation is measured after 4 hours in BθOM T cell hybridoma cells. The data are representative of two independent experiments.

**Supplementary Fig. 9:**
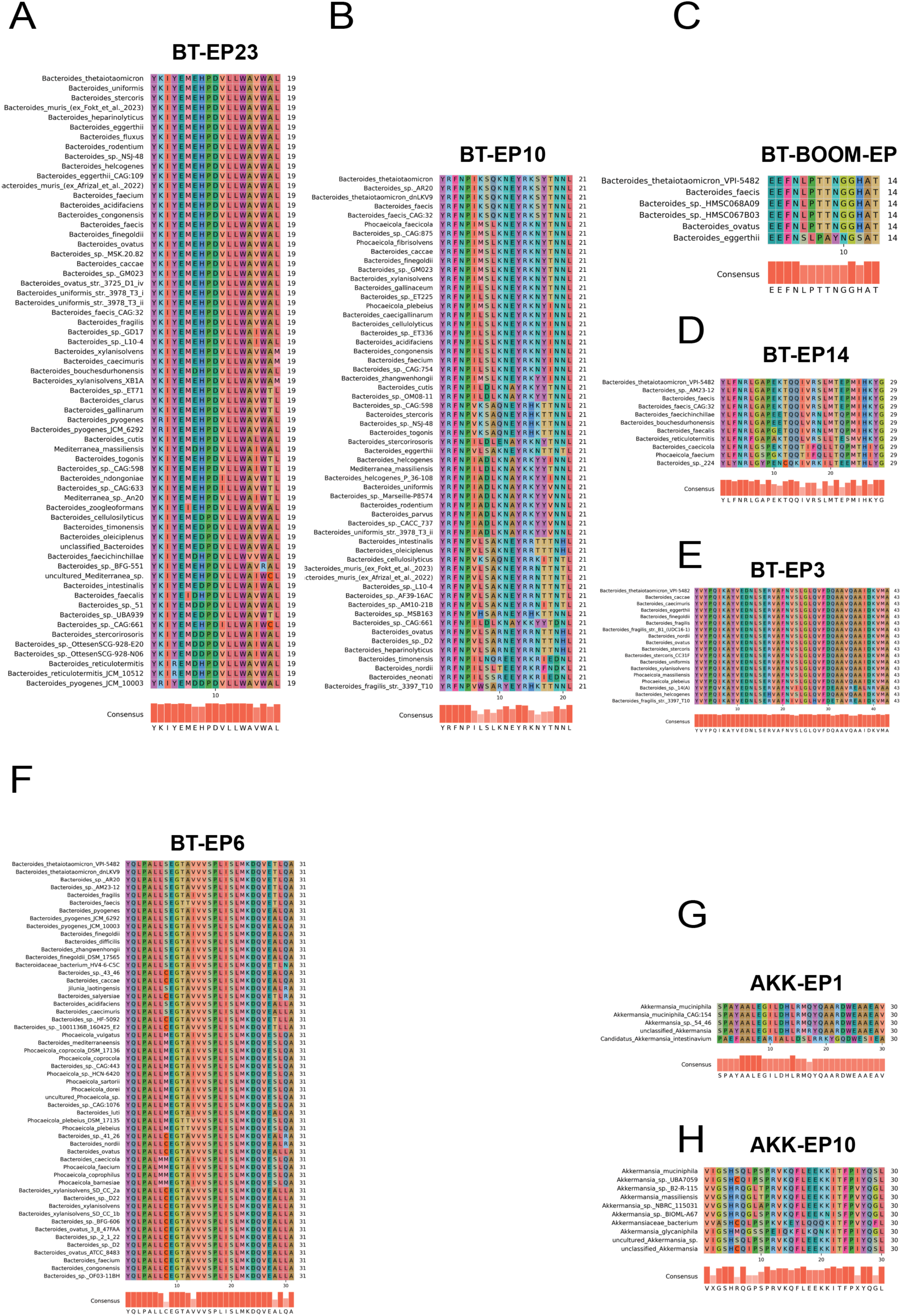
Conservation of variety of BT- and AKK-derived peptides. **(a,b,c,d,e,f)** Multiple sequence alignment (MSA) of BT-EP23, BT-EP10, BT-BθOM-EP, BT-EP14, BT-EP3, and BT-EP6 in the Bacteroides genus of the Bacteroidetes phylum of bacteria. (g,h) MSA of AKK-EP1 and AKK- EP10 in the Akkermansia genus of the phylum Verrucomicrobiota of bacteria.

## Notes

### Competing Interest Statement

The authors have declared no competing interest.

